# Potential pandemic risk of circulating swine H1N2 influenza viruses

**DOI:** 10.1101/2024.01.05.574401

**Authors:** Valerie Le Sage, Nicole C. Rockey, Kevin R. McCarthy, Andrea J. French, Meredith J. Shephard, Ryan McBride, Jennifer E. Jones, Sydney G. Walter, Joshua D. Doyle, Lingqing Xu, Dominique J. Barbeau, Shengyang Wang, Sheila A. Frizzell, Lora H. Rigatti, Michael M. Myerburg, James C. Paulson, Anita K. McElroy, Tavis K. Anderson, Amy L. Vincent Baker, Seema S. Lakdawala

## Abstract

Influenza A viruses in swine have considerable genetic diversity and continue to pose a pandemic threat to humans. They were the source of the most recent influenza pandemic, and since 2010, novel swine viruses have spilled over into humans more than 400 times in the United States. Although these zoonotic infections generally result in mild illness with limited onward human transmission, the potential for sustained transmission of an emerging influenza virus between individuals due to lack of population level immunity is of great concern. Compiling the literature on pandemic threat assessment, we established a pipeline to characterize and triage influenza viruses for their pandemic risk and examined the pandemic potential of two widespread swine origin viruses. Our analysis revealed that a panel of human sera collected from healthy adults in 2020 has no cross-reactive neutralizing antibodies against an α-H1 clade strain but do against a γ-H1 clade strain. Swine H1N2 virus from the α-H1 clade (α-swH1N2) replicated efficiently in human airway cultures and exhibited phenotypic signatures similar to the human H1N1 pandemic strain from 2009 (H1N1pdm09). Furthermore, α-swH1N2 was capable of efficient airborne transmission to both naïve ferrets and ferrets with prior seasonal influenza immunity. Ferrets with H1N1pdm09 pre-existing immunity had reduced α-swH1N2 viral shedding from the upper respiratory tract and cleared the infection faster. Despite this, H1N1pdm09-immune ferrets that became infected via the air could still onward transmit α-swH1N2 with an efficiency of 50%. Taken together, these results indicate that this α-swH1N2 strain has a higher pandemic potential, but a moderate level of impact since there is reduced replication fitness in animals with prior immunity.

## Influenza A virus pandemic assessment

Influenza viruses cause acute respiratory infections in humans, and their wide host range provides many sources of strains with human pandemic potential. Influenza viruses exhibit strong host species preferences, which limits interspecies transmission, but they can evolve specific traits that allow sustained transmission within a new species. Although the major natural global reservoir of influenza virus is wild aquatic birds^1^, swine are an important natural host and can act as a mixing vessel for reassortment of the eight viral gene segments of influenza A viruses from different host species. For example, the most recent H1N1 influenza virus pandemic from 2009 (H1N1pdm09) emerged from swine following reassortment events^2,3^. Emergence of future pandemic strains is a continuing threat necessitating the monitoring and characterization of currently circulating swine viruses.

Influenza viruses are classified into subtypes based on the antigenicity of the surface viral glycoproteins, hemagglutinin (HA) and neuraminidase (NA). HA and NA are important determinants of virus infectivity, transmissibility, pathogenicity, and host specificity and evolve seasonally due to antigenic drift. In swine, three endemic subtypes predominate: swH1N1, swH1N2, and swH3N2 (Extended Data Fig. 1A), which have roughly equal detections over the last four and a half seasons^4^. In the United States, the H1 classical swine lineage (1A) is divided into clades including α-H1 (1A.1), Β-H1 (1A.2), and γ-H1 (1A.3), while the pre-2009 human seasonal-origin swine lineage (1B) includes the δ-H1 (1B.2) clades^5^. The majority of circulating swine strains distributed across the United States are classified within three genetically and antigenically distinct clades from the H1 1A classical swine lineage (1A.1.1.3, 1A.3.3.2, 1A.3.3.3: Extended Data Fig. 1B, 1C and 2)^4,6^. Since the 2010-2011 influenza season, there have been 18 H1N1, 35 H1N2 and 439 H3N2 infections in humans with variants of swine origin in the United States, with six from the α-H1 clade and 21 from γ-H1 clade (https://gis.cdc.gov/grasp/fluview/Novel_Influenza.html).

The current genetic diversity of influenza A virus (IAV) in swine reflects reassortment between avian-, swine-, and human-origin viruses, resulting in multiple lineages of the eight gene segments that continue to reassort among endemic swine strains. The subsequent antigenic drift of HA and NA while circulating in swine resulted in viruses to which the human population may have little to no immunity^7^. Given the potential threat of such swine influenza viruses to humans, we created a decision tree to guide the characterization and pandemic risk assessment of endemic swine IAV (Figure 1). Using a combination of both *in vitro* and *in vivo* methods, this decision tree capitalizes on the extensive research that has been conducted since the 2009 H1N1 pandemic on the molecular properties that promote efficient airborne transmission of influenza^8–20^. In this study, we assessed the pandemic potential of the γ-H1 (1A.3.3.3) clade strain A/swine/Minnesota/A02245409/2020 (herein referred to as ‘γ-swH1N1’) and an α-H1 (1A.1.1.3) clade strain A/swine/Texas/A02245420/2020 (herein referred to as ‘α-swH1N2’). These swine IAV clades were prioritized based on: detection frequency (Extended Data Fig. 1B); geographical distribution (Extended Data Fig. 1C); reported human variant events; significant loss in cross-reactivity to human seasonal vaccines or pre-pandemic candidate vaccine virus antisera^7^; limited detection by human population sera^7^; and interspecies transmission from pigs to ferrets^21^.

**Figure 1.**
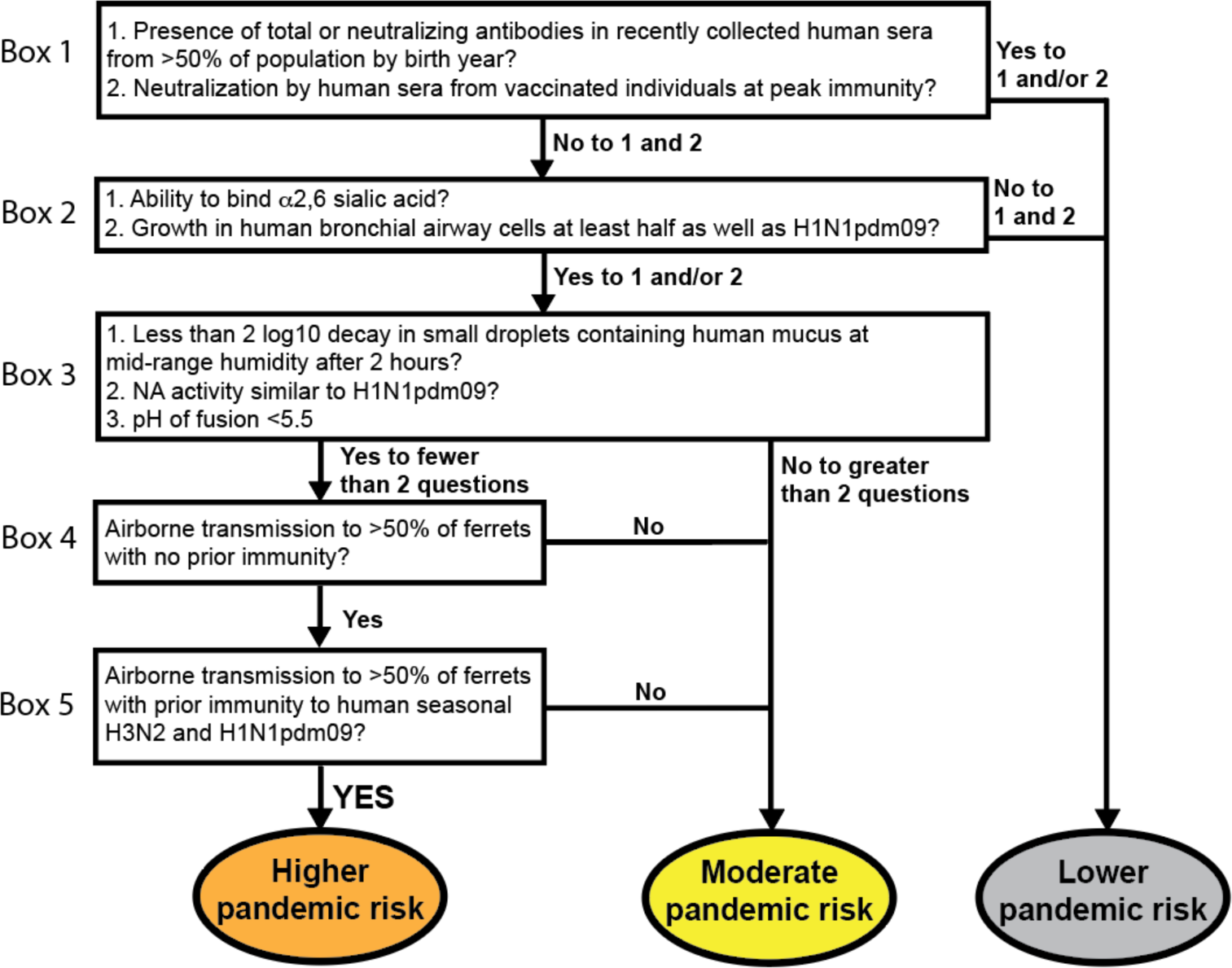
Decision tree of influenza virus pandemic threat assessment.

A pandemic virus represents an antigenic shift, where a large proportion of the population is vulnerable due to a lack of immunity to this novel strain. To assess the presence of cross-reactive influenza virus-specific antibodies (Figure 1, Box 1), human sera collected from healthy adults in Pennsylvania during the fall of 2020 were sorted by birth year and used in hemagglutination inhibition (HAI) (Figure 2A) and neutralization assays (Figure 2B). The prevalence of HAI and/or neutralizing antibodies against γ-swH1N1, α-swH1N2, or H1N1pdm09 was determined, and a threshold HAI titer of 40 was used as it is generally recognized as corresponding to a 50% reduction in the risk of infection^22,23^. H1N1pdm09 and γ-swH1N1-active antibodies as well as neutralizing antibodies were found across all birth year cohorts tested, whereas no HAI titer or neutralizing antibodies were detected against α-swH1N2 in any of the birth years tested (Figure 2A and 2B). In addition, sera from individuals who recently received an influenza virus vaccine were tested to analyze samples with peak immunity from circulating antibodies (Figure 2C). Recently vaccinated individuals had neutralizing antibodies against H1N1pdm09 and γ-swH1N1, but not α-swH1N2 (Figure 2C). Based on the decision tree (Figure 1, Box 1), the presence of cross-reacting antibodies against γ-swH1N1 would funnel the virus to a lower pandemic risk, while α-swH1N2 would require further characterization. However, for this study we proceeded to characterize both γ-swH1N1 and α-swH1N2 to provide empirical evidence for the decision tree criteria.

**Figure 2.**
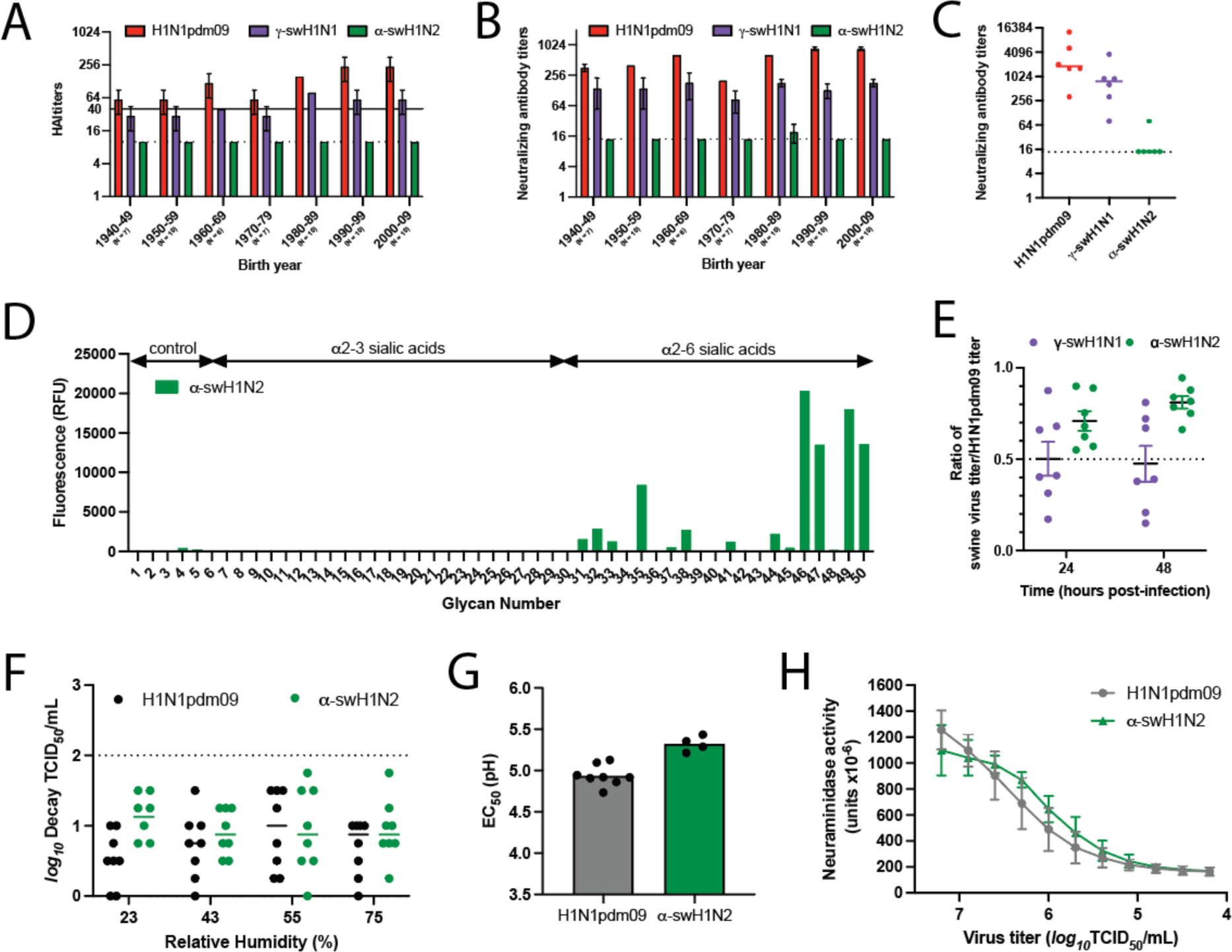
*In vitro* characterization of swine γ-H1N1 and α-H1N2 influenza viruses. Pooled sera from the indicated number of humans for each decade of birth were tested for antibodies to H1N1pdm09 (red bars), γ-swH1N1 (purple bars), and α-swH1N2 (green bars) by HAI **(A)** and neutralization **(B)** assay. **(C)** Sera from individuals vaccinated in October 2021 (14 to 21 days post-vaccination) were assessed for cross-reactive neutralizing antibodies. Each dot represents an individual. Dashed lines indicate the limit of detection for each assay. Solid line indicates an HAI titer of 40, which corresponds to a 50% reduction in the risk of influenza virus infection. **(D)** Binding of α-swH1N2 virus to a sialoside microarray containing glycans with α2-3 or α2-6 linked sialic acids representing avian-type and human-type influenza receptors, respectively. Bars represent the fluorescence intensity of bound α-swH1N2. Glycan structures corresponding to numbers are shown on the x-axis are found in Extended Data Table 1. **(E)** Replication of swine influenza virus in human bronchial epithelial (HBE) air-liquid interface cell cultures. HBE cell cultures were infected in triplicate with 10^3^ TCID_50_ (tissue culture infectious dose 50) per well of H1N1pdm09, γ-swH1N1, or α-swH1N2. The apical supernatant was collected at the indicated time points and virus titers were determined on MDCK cells using TCID_50_ assays. A ratio of swine virus titer relative to H1N1pdm09 titer at 24 and 48 hours of all HBE patient cell cultures is shown. Each dot represents an average of three transwells from seven different HBE patient cultures. **(F)** Stability of α-swH1N2 influenza virus in small droplets over a range of relative humidity (RH) conditions. Aqueous saturated salts were placed at the bottom of a glass desiccator, which was monitored over the duration of the experiment using an Onset HOBO temperature/RH logger. Ten 1uL droplets of pooled virus from panel E were spotted into the wells of a tissue culture dish for 2 hours. Decay of the virus at each RH was calculated compared to the titer of ten 1uL droplets deposited and immediately recovered from a tissue culture dish. Log_10_ decay of HBE-propagated H1N1pdm09 (black) and α-swH1N2 (green) is shown and represents mean values ± standard deviations from three biological replicates performed in triplicate. **(G)** H1N1pdm09 (grey) and α-swH1N2 (green) viruses were incubated in PBS solutions of different pHs for 1 hour at 37°C. Virus titers were determined by TCID_50_ assay and the EC_50_ values were plotted using regression analysis of the dose-response curve. The reported mean (±SD) corresponds to four biological replicates, each performed in triplicate. **(H)** The NA activities of H1N1pdm09 (grey) and α-swH1N2 (green) were determined using an enzyme-linked lectin assay with fetuin as the substrate. Viruses were normalized for equal infectivity and displayed data are the mean (±SD) of three independent experiments performed in duplicate.

## Molecular characterization of swine strains

The H1N1pdm09 HA segment is of swine-origin from the classical H1 lineage^3^. To examine the similarities between the three strains, amino acid differences of the γ-swH1N1 (Extended Data Fig. 3A) and the α-swH1N2 (Extended Data Fig. 3C) HA were mapped onto the H1N1pdm09 HA structure. The γ-swH1N1 HA has 46 amino acid differences as compared to the H1N1pdm09 HA, while α-swH1N2 has 86. Similarity between γ-swH1N1 and H1N1pdm09 HA likely accounts for the cross-neutralizing and cross-receptor blocking antibodies present in human serum (Figure 2 and Extended Data Fig. 3A, purple residues). Diversity in the α-swH1N2 HA is greatest in the globular HA head domain, at sites surrounding the receptor binding site (RBS) (130-strand, 140-loop, 150-loop, 190-helix and the 220-loop^24^) (Extended Data Fig. 3C, green residues). The otherwise conserved RBS is responsible for engaging cell surface sialic acids (SA). In the 130-strand, α-swH1N2 has a two-amino acid deletion (Extended Data Fig. 3D, yellow residues), which may impact antibody binding. Additionally, γ-swH1N1 and α-swH1N2 have an additional putative glycosylation site at the same position on the side of the HA head domain, whereas α-swH1N2 has a second putative glycosylation site near the apex of the HA and its three-fold axis of symmetry (Extended Data Fig. 3A and 3C, pink residues). The evolution of glycosylation sites is thought to contribute to immune escape by shielding antigenic sites on HA^25–27^. During the H1N1 2009 pandemic, differences in the number of putative glycosylation sites between H1N1pdm09 and seasonal viruses were associated with the lack of cross-neutralizing antibodies^28^. Differences in amino acids and glycosylation sites in the α-swH1N2 HA head could contribute to the lack of detectable cross-reactive antibodies observed in Figure 2 and Extended Data Table 1 compared to the γ-swH1N1 or alter receptor preference.

Receptor preference of influenza A viruses is a critical host adaptive property and one known to be important for successful adaptation of influenza viruses to the human population^29^. Human and swine influenza viruses are known to have an α2-6 SA preference, while avian influenza viruses have an α2-3 SA preference. Analysis of H1N1pdm09 and recently circulating human seasonal H3N2 viruses suggests that human viruses adapt to preferential recognition of extended glycans capped with α2-6 SA^29–33^. Analysis of α-swH1N2 receptor specificity using a glycan array with a focused panel of α2-3- and α2-6-linked sialoside glycans showed a strict specificity for glycans with α2-6 sialic acids. For N-linked glycans extended with 1-3 LacNAc (Galβ1-4GlcNAc) repeats, clear preference is shown for extended glycans with two (#47, #49) or three (#47, #50) LacNAc repeats over those with a single LacNAc repeat (#45, #48) (Figure 2D and Extended Data Table 1). Thus, the α-swH1N2 virus exhibits a receptor specificity well adapted for human-type receptors.

To assess fitness of swine viruses to replicate within the human respiratory tract, replication capacity of γ-swH1N1 and α-swH1N2 was determined in human bronchial epithelial (HBE) patient cell cultures grown at an air-liquid interface (Figure 2E). Multiple human HBE cultures were tested, and an H1N1pdm09 virus control was included in all experiments. The ratio of swine virus titer over H1N1pdm09 virus titer for each HBE culture is reported. The representative γ-swH1N1 strain replicated approximately half as well as H1N1pdm09, whereas the representative α-swH1N2 strain had a titer ratio of 0.71 and 0.81 at 24 and 48 hours, respectively (Figure 2E). These data indicate that, regardless of deletions in the RBS 130-loop (Extended Data Fig. 3D, yellow residues), α-swH1N2 replicates to levels similar to H1N1pdm09 (Figure 1, Box 2) and would support α-swH1N2 being selected for additional characterization of parameters correlated with efficient human-to-human transmission of influenza viruses (Figure 1, Box 3).

Airborne transmission requires viral persistence in expelled aerosols and droplets, which can be influenced by environmental conditions, including relative humidity (RH)^34^ or respiratory secretions like HBE airway surface liquid^35,36^. To study the impact of RH on influenza virus viability, droplets of H1N1pdm09 and α-swH1N2 viruses propagated from HBE cultures in Figure 2E were exposed to different RH conditions. HBE-propagated H1N1pdm09 and α-swH1N2 experienced very little decay in infectivity at all RH tested (Figure 2F). These data indicate that α-swH1N2 expelled in small droplets in the presence of human respiratory secretions remains viable over a range of RH conditions, which is important for efficient airborne transmission and viral persistence.

In addition to receptor binding, HA-mediated membrane fusion between the viral envelope and cellular endosome is required for viral entry and is driven by pH changes. A conformational change in the HA from human influenza viruses is triggered between pH 5.3 and 5.5, while avian HA proteins are triggered at a higher pH range of 5.5 to 6.2, suggesting that human adaptation necessitates increased acid stability^37^. To determine the pH at which HA undergoes its conformational change, an acid stability assay was performed on H1N1pdm09 and α-swH1N2, as a surrogate for the pH of fusion^20,38^. The pH that reduces the viral titer by 50% (EC_50_) for α-swH1N2 was 5.3, which was similar to H1N1pdm09 at 5.0 (Figure 2G), indicating that α-swH1N2 has a pH of fusion comparable to human influenza viruses, which is below pH 5.5.

The neuraminidase activity of the NA receptor is necessary to cleave SA from the host cell surface and release the virus. A functional balance between HA and NA is necessary for airborne transmission of swine viruses^18,19,39^. Higher NA activity has also been implicated in the efficient airborne transmission of H1N1pdm09 compared to its swine precursor strains, which had very little NA activity^12^. To measure NA activity, we used an enzyme-linked lectin assay with fetuin as a substrate and a bacterial neuraminidase standard. The NA activity of α-swH1N2 was observed to be similar to that of H1N1pdm09 (Figure 2H). Taken together, these *in vitro* results indicate that α-swH1N2 has the molecular features consistent with a virus capable of airborne transmission and requires further characterization.

## Swine α-H1N2 airborne transmission in ferrets

Following the decision tree criteria (Figure 1), we next characterized α-swH1N2 *in vivo* for the efficiency of airborne transmission in the ferret model (Figure 1, Box 4). Epidemiologically successful human seasonal influenza viruses transmit to naïve recipients after a 2-day exposure^40^. Using this methodology, experimentally infected α-swH1N2 donors were housed with naïve recipients in cages where the animals were separated by a divider. A successful transmission event was defined as recovery of infectious virus in recipient nasal secretions or seroconversion at 21 days post-infection (dpi). In the infected donors, α-swH1N2 was detected in nasal secretions on 1, 2, 3 and 5 dpi (Figure 3A, green bars). Four of four recipients without prior immunity shed α-swH1N2 starting 2 days post-exposure (dpe) (Figure 3A, gray bars). All recipient animals seroconverted at 14 dpi, with increases in antibody titers the following week (Extended Data Table 2). These data indicate that α-swH1N2 transmits efficiently to animals without prior immunity within a short 2-day exposure, similar to published reports of H1N1pdm09^40^.

**Figure 3.**
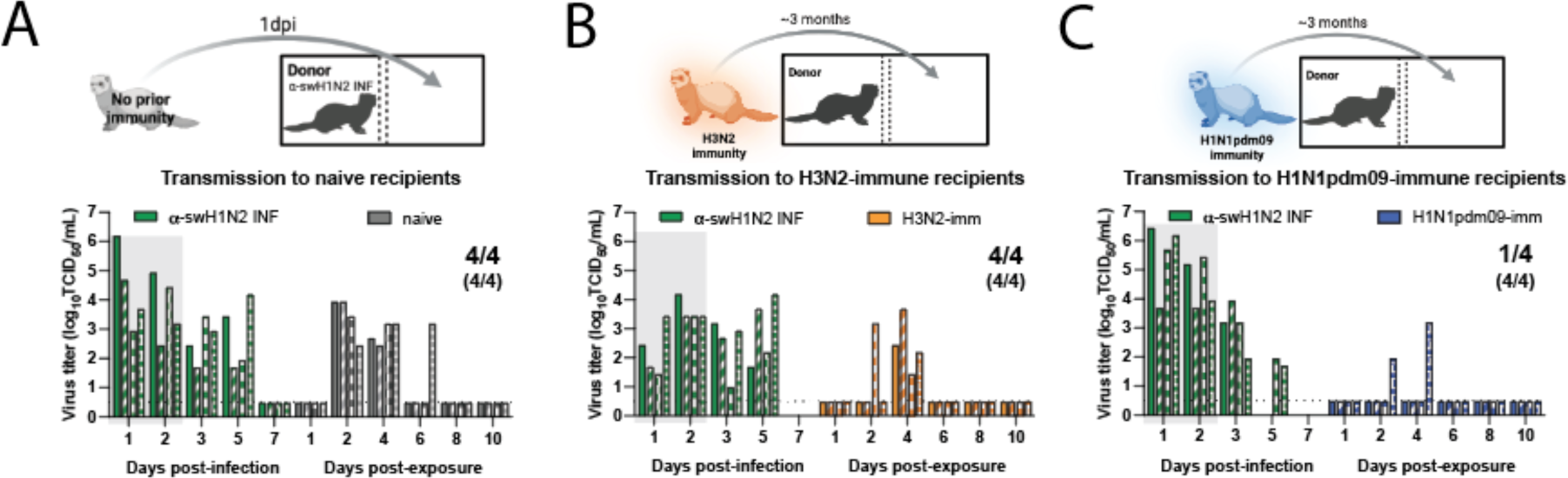
Swine α-H1N2 transmits efficiently via the air after a short exposure. **(A)** Schematic of experimental procedure to naïve recipients. Shaded gray box depicts exposure period. Four donor ferrets were infected intranasally with α-swH1N2 (α-swH1N2 INF), as in Methods. Recipient ferrets with no prior immunity (naïve recipients) were placed in the adjacent cages at 24 hours post-infection for two continuous days. **(B)** Schematic of procedure, whereby four ferrets were infected with H3N2 A/Perth/16/2009 strain (H3N2-imm) 137 days prior to acting as recipients to α-swH1N2 infected donors. Four donor ferrets were infected with α-swH1N2 and H3N2-imm recipients were placed in the adjacent cage 24 hours later. **(C)** Schematic of procedure, whereby four ferrets were infected with H1N1pdm09 (H1N1pdm09-imm) 126 days prior to acting as recipients to α-swH1N2 infected donors. H1N1pdm09-imm recipients were placed in the adjacent cage 24 hours later. Nasal washes were collected from all ferrets on the indicated days and titered for virus by TCID_50_ (tissue culture infectious dose 50). Each bar indicates an individual ferret. For all graphs, the number of recipient ferrets with detectable virus in nasal secretions out of four total is shown; the number of recipient animals that seroconverted at 14- or 21-days post α-swH1N2 exposure out of four total is shown in parentheses. Gray shaded box indicates shedding of the donor during the exposure period. The limit of detection is indicated by the dashed line.

Pandemic influenza viruses do not emerge in immunologically naïve populations as most individuals have experienced influenza by the age of 5^41^. We have previously established a pre-immune ferret model that can be used to assess the pandemic potential of emerging strains in the context of prior immunity^40^. To determine the impact of pre-existing immunity on the transmission efficiency of α-swH1N2, four recipient ferrets were first infected with the H3N2 A/Perth/16/2009 strain (‘H3N2-imm recipient’) or H1N1pdm09 (‘H1N1pdm09-imm recipient’). Roughly 4 months later, once the response to the primary infection was allowed to wane^40,42–45^, these ferrets were then exposed to infected α-swH1N2 donors for 2 days (Figure 3B and 3C). In replicate 1, four of four H3N2-imm recipients shed α-swH1N2 at 4 dpe (Figure 3B), whereas only two of four shed virus in replicate 2 (Extended Data Fig 4). All H3N2-imm recipients that shed virus also seroconverted with increasing antibody titers over time (Extended Data Table 2). All four of four H1N1pdm09-imm recipients seroconverted at 13dpe and had rising antibody titers at 20 dpe (Extended Data Table 2). Intriguingly, only one of four H1N1pdm09-imm recipients shed detectable levels of α-swH1N2 (Figure 3C). It is possible that shedding of virus was missed in the recipients either because the nasal wash samples were not taken at 3 dpe or that replication of the virus was occurring in a place that was not sampled by the nasal wash, such as the mid-turbinates, nasopharynx, trachea, or lungs. However, based on serology we can conclude that all four H1N1pdm09 pre-immune animals were infected (Extended Data Table 2). These data suggest that α-swH1N2 can transmit to animals with prior immunity, which categorizes α-swH1N2 into the higher pandemic risk. However, whether naturally infected ferrets with prior immunity could spread the virus onward is unclear.

## Potential α-swH1N2 pandemic severity

Person-to-person airborne transmission is a concern for pandemic emergence and can be experimentally assessed using transmission chain experiments. Given the lack of detectable shedding of α-swH1N2 in three of four H1N1pdm09-imm recipients yet seroconversion in all four recipients in Figure 3C, we examined whether H1N1pdm09-imm recipients would shed enough virus to onward transmit α-swH1N2 to naïve recipients. Two independent replicate transmission chains were performed with four α-swH1N2 infected donors being exposed to H1N1pdm09-imm recipients in the adjacent cage for 2 days. The exposed H1N1pdm09-imm recipients (R1) were then transferred to a new cage to act as donors to naïve recipients (R2) (Figure 4A). In the first replicate (Figure 4B), two of four R1 ferrets shed α-swH1N2, whereas in the second replicate (Figure 4C), all four R1 ferrets had α-swH1N2 in their nasal secretions. Much of the viral shedding was observed on day 3 post exposure, which may account for the absence of robust shedding in Figure 3C. When R1 ferrets became donors (Figure 4B and 4C, pink box), only 50% of the infected donors transmitted α-swH1N2 onward to influenza immunologically naïve recipients. To further examine the ability of α-swH1N2 to transmit from H1N1pdm09-imm ferrets in the context of pre-existing immunity, a transmission chain experiment was performed using R2 recipients with H1N1pdm09 immunity (Figure 4D). To ensure that the exposure window of viral shedding of R1 was captured, rapid antigen tests were performed immediately following the nasal wash, when positive that animal was transferred into a cage to act as a donor animal to an H1N1pdm09 R2 immune ferret (Figure 4D). In this study, all three R1 became infected and of these three α-swH1N2-infected H1N1pdm09-imm R1 recipients only one onward transmitted to the H1N1pdm09-imm R2 recipients (Figure 4D). Seroconversion only occurred in R1 and R2 recipients that shed detectable α-swH1N2 virus (Extended Data Table 2). These data suggest that onward transmission of α-swH1N2 is possible, even in the context of pre-existing immunity, contributing to a higher risk potential of α-swH1N2.

**Figure 4.**
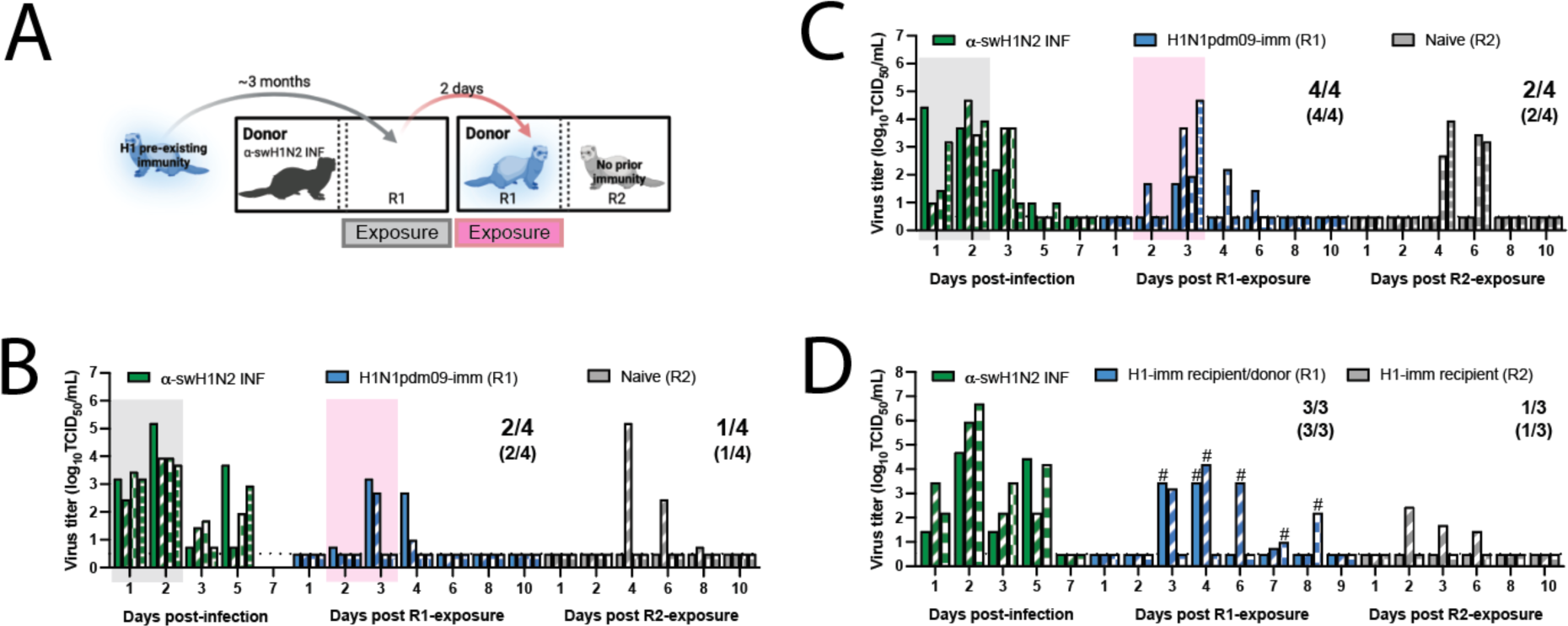
Swine H1N2 virus transmission chain. **(A)** Schematic of transmission chain experiment in panels B and C. Four ferrets were infected with H1N1pdm09 (R1) 127 days before being exposed to a donor that was infected with α-swH1N2 24 hours prior. After a 2-day exposure, H1N1pdm09-imm recipients were transferred to a new transmission cage to act as the donors. The transfer of each R1 animal was done without knowledge of its infection status. A naïve recipient (R2) was immediately placed in the adjacent cage and exposed for 2 days. The transmission chain experiment was performed two independent times. Nasal secretions were collected for all animals on the indicated days post-infection or post-exposure, with each bar representing the virus titer shed by an individual animal for replicate 1 **(B)** and replicate 2 **(C)**. Gray shaded boxes indicate the days upon which the α-swH1N2 infected (α-swH1N2 INF) donor was exposing the H1N1pdm09-imm recipient (R1), and the pink shaded box indicates the days upon which R1 was acting as the donor to R2. Limit of detection is denoted by a dashed line. The numbers in parentheses indicate the proportion of animals that seroconverted. **(D)** Donors were infected with α-swH1N2 24 hours prior to exposing H1N1pdm09-imm recipients. Nasal washes from R1 recipients were collected and immediately tested using a rapid antigen test. Once positive for influenza virus antigen, the H1N1pdm09-imm R1 recipient was moved into a new transmission cage to act as the donor and expose an H1N1pdm09-imm R2 recipient for 2 days. # indicates the 2-day window in which each of the R1 ferret exposed the R2 recipients.

We next examined the impact of prior influenza virus exposure on α-swH1N2 replication and pathogenesis; ferrets with no prior immunity, pre-existing immunity against H3N2 (H3N2-imm) or H1N1pdm09 (H1N1pdm09-imm) were intranasally infected with α-swH1N2 and their nasal secretions were collected over time. No difference in α-swH1N2 titers was observed between H3N2-imm ferrets and those with no prior immunity from 1-3 dpi, however, H3N2-imm ferrets cleared α-swH1N2 by 5 dpi (Figure 5A). This observation is consistent with our previous reports of a reduced viral shedding period in animals with heterosubtypic immunity^40,46^. Interestingly, ferrets with pre-existing H1N1pdm09-imm shed significantly less α-swH1N2 virus on 1, 2, and 3 dpi as compared to ferrets with no prior immunity (Figure 5B).

**Figure 5.**
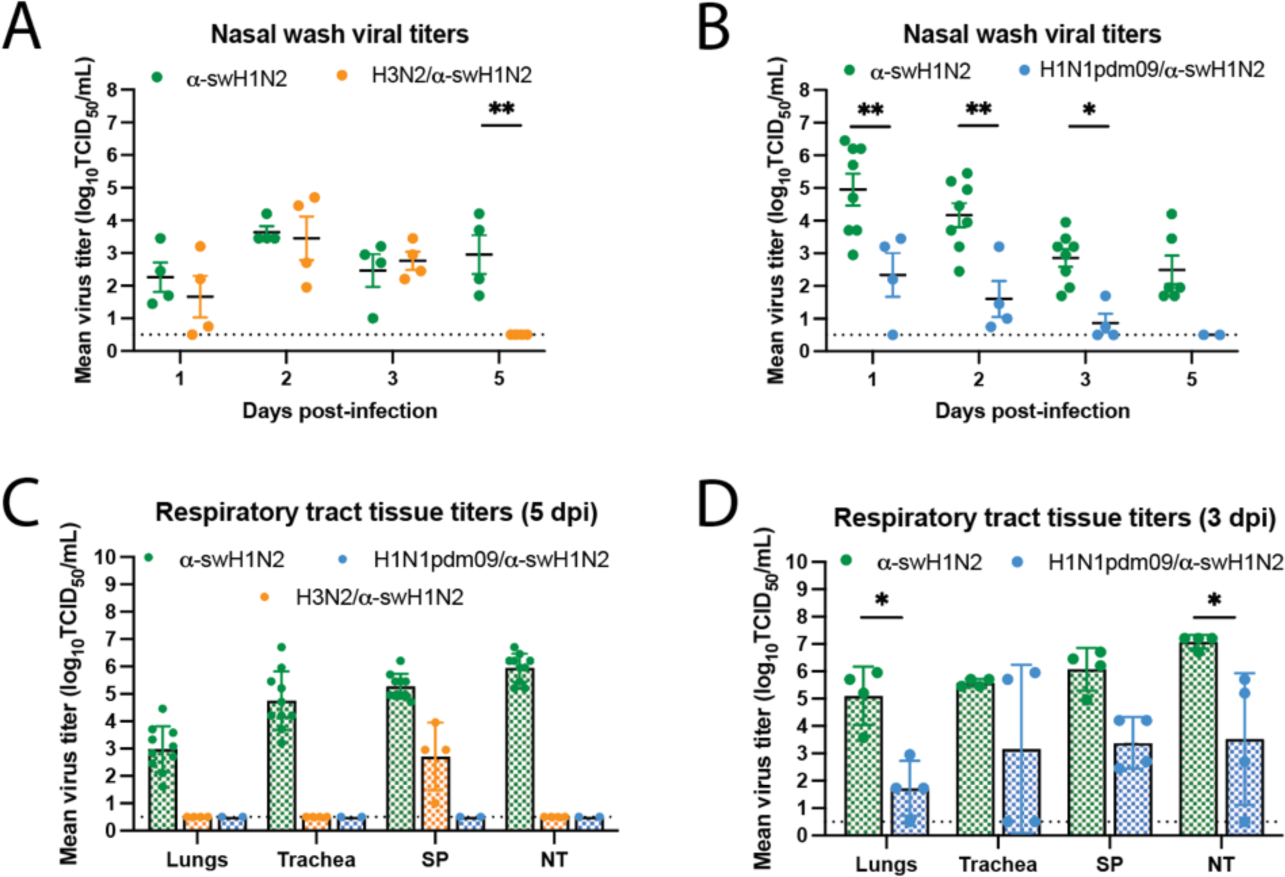
H1N1pdm09-immune ferrets have reduced α-swH1N2 viral titers in nasal secretions and tissues. **(A)** Ferrets with no pre-existing immunity (α-swH1N2, N=4) or those infected with H3N2 137 days prior (H3N2/α-swH1N2, N=4) were intranasally infected with α-swH1N2. The mean ± SD viral titers from nasal secretions are shown with each circle representing an individual animal. Two-way ANOVA analysis was used to determine statistically significant differences (** p<0.005). **(B)** α-swH1N2 mean ± SD viral titers from nasal secretions from animals with no prior immunity (α-swH1N2 INF, N=8) or those infected with H1N1pdm09 126 days prior (H1N1pdm09/α-swH1N2, N=4). Two-way ANOVA analysis was used to determine statistically significant differences (** p<0.005 and * p<0.05). **(C)** Respiratory tissues from α-swH1N2 infected ferrets with no prior immunity (green; N=10), H3N2 prior immunity (orange; N=4), or H1N1pdm09 prior immunity (blue; N=2) were collected at 5 dpi. Graphs show the mean ± SD viral titers. SP-soft palate, NT-nasal turbinates. **(D)** Respiratory tissues from α-swH1N2 infected ferrets with no prior immunity (green; N=4) or H1N1pdm09 prior immunity (blue; N=4) were collected at 3 dpi. Graphs show the mean ± SD viral titers. Two-way ANOVA analysis was used to determine statistically significant differences (* p<0.05). The dashed line indicates the limit of detection for all graphs.

To characterize tissue-specific α-swH1N2 replication, infected ferrets from panels 5A and 5B were sacrificed at 5 dpi and the respiratory tract was collected for viral titration (Figure 5C). In ferrets with no prior immunity, robust replication was detected in the lungs, trachea, soft palate, and nasal turbinates, whereas H3N2-imm ferrets only had detectable α-swH1N2 in the soft palate (Figure 5C). H1N1pdm09-imm ferrets had completely cleared α-swH1N2 from their respiratory tracts on day 5, as no detectable infectious virus was detected in any of the collected tissues (Figure 5C). Since viral titers from nasal washes were already reduced in these animals by 3 dpi, we next assessed viral replication in the respiratory tract at this time point. H1N1pdm09-imm and non-immune ferrets were infected with α-swH1N2 and sacrificed on day 3 (Figure 5D). H1N1pdm09-imm ferrets had detectable α-swH1N2 in the respiratory tract, although viral titers were significantly less in the lungs and nasal turbinates compared to animals without prior immunity. Taken together, these data indicate that prior H1N1pdm09 immunity can reduce the viral load in the ferret respiratory tract and decrease time to clearance of α-swH1N2.

To extend the observation on viral titers, we compared the lung pathology of α-swH1N2-infected ferrets with no prior immunity to those with H1N1pdm09 pre-existing immunity (from Figure 5D). Regardless of immunity, at 3 dpi α-swH1N2-infected ferrets had bronchial glands that were multifocally necrotic and contained neutrophils as well as peripheral lymphocytes, whereas uninfected ferrets had intact glands and no inflammation (Extended Data Fig. 5). The bronchioles from infected ferrets with no prior immunity were ulcerated and had evidence of macrophage and neutrophil accumulation within the airway lumen, whereas, H1N1pdm09-imm ferrets were similar to uninfected ferrets in that their bronchioles were clear of cellular debris with intact ciliated columnar lining epithelium (Extended Data Fig. 5). Furthermore, H1N1pdm09-imm alveolar interstitium had large airways that were clear with mild to moderate peripheral lymphocytic infiltrates and blood vessels that were multifocally surrounded by edema and lymphocytic infiltrates (Extended Data Fig. 5). In the absence of prior immunity, the large airways of the alveolar interstitium were partially ulcerated and filled with immune cells, and the alveolar spaces were filled with fibrin edema (Extended Data Fig. 5). These data indicate that pre-existing H1N1pdm09 immunity can reduce the pathology caused by α-swH1N2 infection.

Lastly, we examined the clinical outcomes for α-swH1N2 infected ferrets during these studies by cataloging the activity, weight loss, and other signs of the animals^47^. Intranasally α-swH1N2-infected ferrets with no prior immunity and H3N2-imm displayed a similar number of symptoms, while intranasally infected H1N1pdm09-imm ferrets displayed almost no symptoms (Extended Data Fig. 6A). Ferrets with no prior immunity displayed a greater range of symptoms than those with pre-existing immunity (Extended Data Fig. 6C). In airborne-infected animals, all ferrets, regardless of immunity, displayed a similar mean total number of symptoms (Extended Data Fig. 6B), which included similar clinical signs over multiple days being reduced activity scores, nasal discharge and weight loss (Extended Data Fig. 6D). Overall, ferrets intranasally or airborne infected with α-swH1N2 had mild symptoms, which varied by the category of symptoms.

## Discussion

Identification of emerging respiratory viruses with pandemic potential is critical for enacting preparedness measures to mitigate their impact. Swine viruses are particularly concerning, given their agricultural importance that places them within close physical proximity to humans and the wide diversity of swine influenza strains^48^. Current risk assessment of pandemic threats is done through the WHO and CDC risk assessment tools^49,50^, which use subject-area expert opinion to assign weighted scores for various categories and limited experimental data derived from multiple different *in vitro* and *in vivo* sources. In this study, we present a streamlined, adaptable strategy to experimentally triage influenza viruses that reduces the need for complete virus characterization since certain criteria must be met before proceeding to the next box in the decision tree. This pipeline represents a breathable framework that can and will be updated as additional data from characterization studies are conducted.

Using our decision tree, we analyzed representative circulating swine H1 strains from the alpha and gamma genetic clades that have a wide geographic distribution, are frequently detected in swine populations in the United States (Extended Data Fig. 1C), and have exhibited sporadic human spillover events^51^. Previous representatives of the α-swH1N2 clade were shown to have antigenic distance from human vaccine strains, reduced recognition by human sera from two different cohorts^6^, and transmitted efficiently from infected pigs to naive recipient ferrets^52^. While highly efficient at controlling antigenically similar influenza viruses, antibodies directed towards HA become less effective over each subsequent flu season as surface glycoproteins rapidly mutate through antigenic drift. No cross-neutralizing antibodies were detected against α-swH1N2 in H1N1pdm09- or H3N2-imm ferrets (Extended Data Table 2), suggesting that an initial infection with human seasonal viruses does not produce antibodies that cross-neutralize, and this was consistent with our human serum data (Figure 2). Interestingly, human sera across all birth years tested had variable levels of anti-N2 antibodies (Extended Data Fig. 7), which may suggest that this NA-based immunity could provide some level of protection in a subset of the population^53–56^. Our prior work previously determined that prior immunity can influence the susceptibility to heterosubtypic viruses in a mechanism not mediated by neutralizing antibodies^40^. Thus, prior immunity from divergent strains can impact susceptibility of viruses through the air. We found that α-swH1N2 transmitted efficiently through the air to ferrets regardless of immune status, but the severity of disease after experimental infection with α-swH1N2 was lower in animals with prior immunity. A similar phenomenon may explain the lower-than-expected morbidity and mortality of the 2009 pandemic in humans^57^.

Protection against emerging influenza virus strains in hosts without neutralizing antibodies can be conferred from CD8^+^ T cells, which recognize conserved internal influenza virus proteins. Although prior adaptive immunity may not prevent influenza virus infection, CD8^+^ T cells that display cross-reactivity against different subtypes of influenza virus have been linked to more efficient clearance of virus and faster recovery from illness^58–60^. Indeed, prior immunity to human seasonal viruses was not protective against α-swH1N2 airborne infection (Figure 3C and D). Encouragingly, experimentally infected ferrets with pre-existing immunity were able to clear α-swH1N2 faster and H1N1pdm09 immunity resulted in an overall decrease in virus shedding over time (Figure 5B) and decreased lung pathology early during infection (Extended Data Fig. 5). However, the lack of disease severity in immune animals may also provide an opportunity for this virus to spread undetected and gain a foothold in the population, creating a pandemic risk. Taken together, our data demonstrate that this α-swH1N2 virus strain poses a high pandemic risk that warrants continued surveillance efforts to capture zoonotic events and an increased campaign to vaccinate swine against this H1 clade to reduce the amount of virus in source populations.

## Acknowledgements

This project has been funded in part with Federal funds from the National Institute of Allergy and Infectious Diseases, National Institutes of Health, Department of Health and Human Services, under Contract No. 75N93021C00015; the United States Department of Agriculture, Agricultural Research Service, project number 5030-32000-231-000-D, Cystic Fibrosis Foundation Research Development Program to the University of Pittsburgh, and Burroughs Wellcome CAMS 1013362.02 to AKM. We thank Dr. Daniel Perez for generously providing plasmids. We thank Dr. Rachel Duron for critical review and feedback. The funders had no role in study design, data collection and interpretation, or the decision to submit the work for publication. Mention of trade names or commercial products in this article is solely for the purpose of providing specific information and does not imply recommendation or endorsement by the USDA. USDA is an equal opportunity provider and employer.

## Author Contributions

VL and SSL designed the experiments, analyzed, interpreted the data and wrote the manuscript. VL, NRC, KRM, AJF, MJS, RM, JEJ, SGW and LHR performed the experiments. JDD, LX, DJB, SW, SAF, MMM, JCP, AKM, TKA and ALVB contributed resources and analysis. All authors edited and approved the manuscript.

## Author Information

The authors to declare no competing financial and/or non-financial interests in relation to the work described.

## METHODS

### Genetic analysis and strain selection

All available swine H1 HA sequences from the USA collected between January 2019 and December 2021 were downloaded from the Bacterial and Viral Bioinformatics Research Center (BV-BRC)^61^. These data (n=2144) were aligned with the World Health Organization (WHO)-recommended human seasonal H1 HA vaccine sequences and candidate vaccine sequences. The swine and human IAV HA sequences were aligned using mafft v7^62^, and a maximum likelihood phylogeny for the alignment was inferred, following automatic model selection, using IQ-TREE v2^63^ and visualized using smot v1.0.0^64^ (Extended Data Fig. 2). The evolutionary lineage and genetic clade of each swine HA gene was identified using the BV-BRC Subspecies Classification tool, and the predominant clades and their geographic distribution were identified^4,65^. The 1A and 1B lineages were detected in the USA and the genetic clades 1A.3.3.3 (38%, n=806), 1B.2.1 (29%, n=622), 1A.3.3.2 (12%, n=263), 1A.1.1.3 (11%, n=233), 1B.2.2.1 (5%, n=109) and 1B.2.2.2 (3%, n=69) represented 98% of detections. Given human variant detections, evidence for interspecies transmission, a significant reduction in cross-reactivity to human seasonal vaccines or candidate vaccine viruses, and limited detection by human population sera, we prioritized the 1A.1.1.3 and 1A.3.3.3 clades for characterization^7,21^. Representative selections within these clades were identified by generating an HA1 consensus sequence and identifying the best-matched field strain to the consensus that had an NA and internal gene constellation that reflected the predominant evolutionary lineages detected in surveillance (1A.1.1.3/α-swH1N2, A/swine/Texas/A02245420/2020, 97.2% to HA1 consensus: and 1A.3.3.3/γ-swH1N1, A/swine/Minnesota/A02245409/2020, 98.5% to consensus). Furthermore, the selected viruses had NA and internal gene patterns that matched the predominant evolutionary lineages detected between 2019-2021 (https://flu-crew.org/octoflushow/). The 1A.1.1.3/α-swH1N2 was paired with a N2-2002A gene with a TTTTPT internal gene constellation, and the 1A.3.3.3/γ-swH1N1 was paired with a N1-Classical gene with a TTTPPT internal gene constellation.

### Cells and viruses

Madin Darby canine kidney (MDCK) epithelial cells (ATCC, CCL-34) were maintained in Eagle’s minimal essential medium supplemented with 10% fetal bovine serum, penicillin/streptomycin and L-glutamine. Primary HBE cell cultures were derived from human lung tissue that were differentiated and cultured at an air-liquid interface using a protocol approved by the relevant institutional review board at the University of Pittsburgh^66^. The influenza A virus strains, A/swine/Texas/A02245420/2020 (α-swH1N2, 1A.1.1.3) and A/swine/Minnesota/A02245409/2020 (γ-swH1N1, 1A.3.3.3) were obtained from the National Veterinary Services Laboratories (NVSL) repository for the USDA IAV-S surveillance system. Reverse genetic derived strains of A/California/07/2009 (H1N1pdm09) and A/Perth/16/2009 (H3N2) were a generous gift from Dr Jesse Bloom (Fred Hutch Cancer Research Center, Seattle) and were rescued as previously described in ^67^.

### TCID_50_ assay

MDCK cells were seeded at a density of 10,000 cells per well in 96-well plate three days prior to the assay. Cells were washed with sterile phosphate-buffered saline (PBS) followed by addition of 180 μL of Eagle’s minimal essential medium supplemented with Anti-Anti, L-glutamine and 0.5 μg/mL TPCK-treated trypsin. 20 μL of virus was diluted in the first row and tenfold serial dilutions on cells were performed. The assay was carried out across the plate with the last row as the cell control without virus. The cells were incubated for 96 hours at 37°C in 5% CO2 and scored for cytopathic effect (CPE).

### Serological assays

Hemagglutination inhibition (HAI) was used to assess the presence of receptor-binding antibodies to HA protein from the selected viruses in human sera. Briefly, one-part sera were treated with three parts receptor destroying enzyme (RDE) overnight at 37°C to remove non-specific inhibitors. The following day, the sera was heat inactivated at 56°C for 30 minutes and six parts of normal saline added. In a V-bottom microtiter plate, two-fold serial dilutions of RDE-treated sera were performed and incubated with eight hemagglutinating units of virus for 15 minutes. Turkey red blood cells were added at a concentration of 0.5% and incubated for 30 minutes. The reciprocal of the highest dilution of serum that inhibited hemagglutination was determined to be the HAI titer. The titer of neutralizing antibodies was determined using the microneutralization assay. Human or ferret sera was heat inactivated at 56°C for 30 minutes and serially diluted 2-fold in a 96-well flat-bottom plate. 10^3.3^ TCID_50_ of influenza virus was incubated with the sera for 1 hour at room temperature before being transferred to a 96-well plate on confluent MDCK cells. Sera was maintained for the duration of the experiment and CPE was determined on day 4 post-infection. The neutralizing titer was expressed as the reciprocal of the highest dilution of serum required to completely neutralize the infectivity of 10^3.3^ TCID_50_ of virus on MDCK cells. The concentration of antibody required to neutralize 100 TCID_50_ of virus was calculated based on the neutralizing titer dilution divided by the initial dilution factor, multiplied by the antibody concentration.

### Glycan array

Glycan arrays were prepared as previously described^33,68^. Briefly, glycans were prepared at 100 µM in 150 mM Na3PO4 buffer (pH 8.4) and printed onto NHS-activated glass microscope slides (SlideH, Schott) using a MicroGridII (Digilab) contact microarray printer equipped with Stealth Microarray Pins (SMP3, ArrayIt). Residual NHS was blocked by treatment with 50mM ethanolamine in 50mM borate buffer, pH 9.2 for 1 hour and washed with water. Slides were centrifuged to remove excess water and were stored at −20C. For analysis of receptor specificity glycan arrays were overlayed with culture fluid containing intact influenza virus prepared in MDCK cells for one hour at room temperature. Slides were then washed with phosphate buffered saline (PBS) and water, followed by incubation with biotinylated Galananthus Novalis Lectin (GNL; Vector Labs) at 1ug/mL in 1X PBS for one hour^68^. Sides were washed with PBS and overlayed with 1 µg/ml Streptavidin-AlexaFluor488 (LifeTech) for one hour, and washed with PBS and water. Slides were then scanned using an Innoscan 1100AL microarray scanner (Innopsys). Signal values are calculated from the mean intensities of 4 of 6 replicate spots with the highest and lowest signal omitted and graphed.

### Replication kinetics

Four different HBE patient cell cultures were used (HBE0344, HBE0338, HBE0342, HBE0370). The apical surface of the HBE cells was washed in PBS and 10^3^ TCID_50_ of virus was added per 100 μL of HBE growth medium. After 1 hour incubation, the inoculum was removed and the apical surface was washed three times with PBS. At the indicated time points, 150 μL of HBE medium was added to the apical surface for 10 minutes to capture released virus particles. The experiment was performed in triplicate in at least three different patient cell cultures. Infectious virus was quantified by TCID_50_ using the endpoint method^69^.

### Enzyme-linked lectin assay (ELLA)

The neuraminidase activity was determined using a peanut-agglutinin based ELLA. A 96-well ultra-high binding polystyrene plate was coated with 25 μg/mL of fetuin diluted in coating buffer overnight at 4°C and the excess fetuin was removed using wash buffer (0.01M PBS, pH 7.4, 0.05% Tween 20). Two-fold serial dilutions of 10^7^.^5^ TCID_50_/mL virus stock or 62.5 mU/mL *Clostridium perfringes* neuraminidase (to standardize the viruses between different plates) were performed in a 96-well plate. Serial dilutions were then transferred to the plates coated with fetuin and incubated overnight at 37°C. Plates were thoroughly washed 6 times with wash buffer and incubated in the dark at room temperature with peroxidase-labeled peanut agglutinin solution for 2 hours. O-phenylenediamine dihydrochloride substrate was added for 10 minutes to and the reaction was stopped using sulfuric acid. Absorbance was read at 490 nm. NA activity was assayed in duplicate and performed in three independent replicates.

### pH inactivation assay

The pH of inactivation assay^70^ was used to determine the pH at which HA undergoes its irreversible conformational change. 10 μL of virus stock was incubated in 990 μL of PBS adjusted to the indicated pH values for 1 hour at 37 °C and immediately neutralized by titering on MDCK cells using the TCID_50_ endpoint titration method^69^ to determine the remaining infectious virus titer. The pH that reduced the titer by 50 % (EC_50_) was calculated by regression analysis of the dose-response curves. Each experiment was performed in triplicate in at least three independent biological replicates.

### Stability of stationary droplets

Desiccator chambers containing saturated salt solutions of potassium acetate, potassium carbonate, magnesium nitrate or sodium chloride were equilibrated to 23%, 43%, 55% of 75% relative humidity (RH), respectively. Ten 1 μL droplets of HBE-propagated virus were spotted onto a 6-well plate in duplicate and immediately incubated in the desiccator chamber for 2 hours. Chambers were maintained in a biosafety cabinet for the duration of the experiment and a HOBO UX100011 data logger was used to collect RH and temperature data. After 2 hours, the droplets were collected in 500 μL of L-15 medium, which was titered on MDCK cells using the TCID_50_ endpoint method^69^. Decay was determined by subtracting the titer of the virus aged for 2 hours from the titer of the virus that had been deposited and then immediately recovered.

### Animal ethics statement

Ferret experiments were conducted in a BSL2 facility at the University of Pittsburgh in compliance with the guidelines of the Institutional Animal Care and Use Committee (approved protocol 22061230). Animals were sedated with isoflurane following approved methods for all nasal washes and survival blood draws. Ketamine and xylazine were used for sedation for all terminal procedures, followed by cardiac administration of euthanasia solution. Approved University of Pittsburgh Division of Laboratory Animal Resources (DLAR) staff administered euthanasia at time of sacrifice.

### Human subjects research ethics statement

Human serum samples used in this study were collected from healthy adult donors who provided written informed consent for their samples to be used in infectious disease research. The University of Pittsburgh Institutional Review Board approved this protocol (STUDY20030228). All participants self-reported their age, sex, race, ethnicity, residential zip code, history of travel and immunization. HBE cultures are obtained from deidentified patients (STUDY19100326) and provided from the tissue airway core to for these studies.

### Ferret screening

Four- to six-month-old male ferrets were purchased from Triple F Farms (Sayre, PA, USA). All ferrets were screened by HAI for antibodies against circulating influenza A and B viruses, as described in ‘Serology’ section. The following antigens were obtained through the International Reagent Resource, Influenza Division, WHO Collaborating Center for Surveillance, Epidemiology and Control of Influenza, Centers for Disease Control and Prevention, Atlanta, GA, USA: 2018–2019 WHO Antigen, Influenza A (H3) Control Antigen (A/Singapore/INFIMH-16-0019/2016), BPL-Inactivated, FR-1606; 2014–2015 WHO Antigen, Influenza A (H1N1)pdm09 Control Antigen (A/California/07/2009 NYMC X-179A), BPL-Inactivated, FR-1184; 2018–2019 WHO Antigen, Influenza B Control Antigen, Victoria Lineage (B/Colorado/06/2017), BPL-Inactivated, FR-1607; 2015–2016 WHO Antigen, Influenza B Control Antigen, Yamagata Lineage (B/Phuket/3073/2013), BPL-Inactivated, FR-1403.

### Ferret infections

To generate ferrets with pre-existing immunity against seasonal influenza viruses, ferrets were inoculated intranasally with 10^6^ TCID_50_ in 500 μL (250 μL in each nostril) of recombinant A/California/07/2009 or A/Perth/16/2009. These animals were allowed to recover and housed for 126 to 137 days before acting as a recipient in a transmission experiment or being similarly infected with A/swine/Texas/A02245420/2020.

### Transmission studies

The transmission cage setup was a modified Allentown ferret and rabbit bioisolator cage^12,32^. For each study, four ferrets were anesthetized with isoflurane and inoculated intranasally with 10^6^ TCID_50_ in 500 μL (250 μL in each nostril) of A/swine/Texas/A02245420/2020 to act as donors. Twenty-four hours later, a naïve or immune recipient ferret was placed into the adjacent cage, which is separated by two staggered perforated steel plates welded together one inch apart with directional airflow from the donor to the recipient. Recipients were exposed to the donors for 2 days with nasal washes being collected from each donor and recipient every other day for 11 days. For the transmission chain experiment (Figure 4), after the initial 2-day exposure, the recipients were transferred to the donor side of a new transmission cage where a naïve recipient ferret was on the other side of the divider. These animals were subsequently singled housed following 48 hours. To prevent accidental contact or fomite transmission by investigators, the recipients were handled first and extensive cleaning of gloves, sedation chamber, biosafety cabinet, and temperature monitoring wand was performed between each pair of animals. Sera from donor and recipient ferrets were collected upon completion of experiments to confirm seroconversion. To ensure no accidental contact or fomite transmission during husbandry procedures, recipient animal sections of the cage were cleaned prior to the donor sides, with one cage being done at a time. Fresh scrapers, gloves, and sleeve covers were used for each subsequent cage cleaning. Clinical symptoms such as weight loss and temperature were recorded during each nasal wash procedure and other symptoms such as sneezing, coughing, activity, diarrhea or nasal discharge were noted during any handling events. Once animals reached 10% weight loss, their feed was supplemented with A/D diet twice a day to entice eating. Clinical scoring was previously described in ^47^.

### Tissue collection and processing

The respiratory tissues were collected from euthanized ferrets aseptically in the following order: entire right middle lung, left cranial lung (a portion equivalent to the right middle lung lobe), one inch of trachea cut lengthwise, entire soft palate, and nasal turbinates, as described previously in ^32^. Tissue samples were weighed, and Leibovitz’s L-15 medium was added to make a 10% (lungs) or 5% (trachea) w/v homogenate. Tissues were dissociated using an OMNI GLH homogenizer (OMNI International) and cell debris was removed by centrifugation at 900 xg for 10 minutes. Influenza virus titers were determined by endpoint TCID_50_ assay^69^. The lungs were fixed in 10% neutral buffered formalin and subsequently processed in alcohols for dehydration and embedded in paraffin wax. Sections were stained with haematoxylin and eosin (H&E). The sections were examined ‘blind’ to experimental groups to eliminate observer bias by a board-certified animal pathologist (LHR).

### Data availability

The raw data generated and analyzed during the current study have been archived on FigShare (<u>10.6084/m9.figshare.c.6982905</u>). Phylogenetic analyses and associated strain selection data are archived at https://github.com/flu-crew/datasets

---

**Extended Data Table 2:**
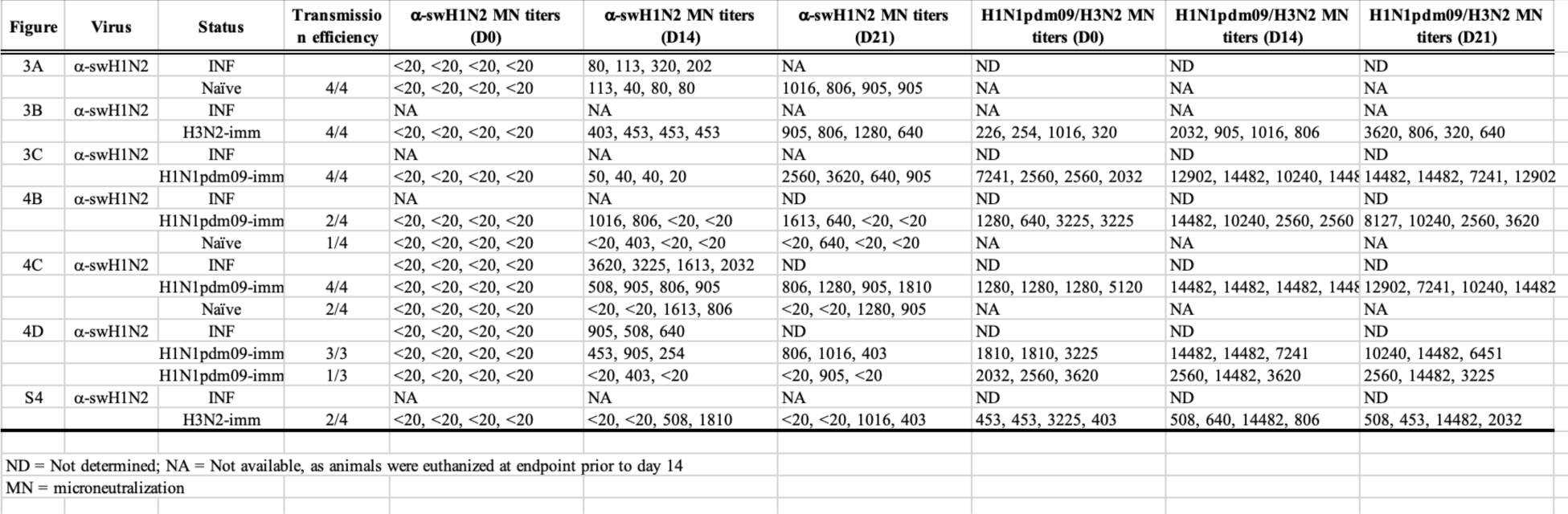
Serology or donor and rtcipient rerrets for each transmission study.

**Extended Data Figure 1.**
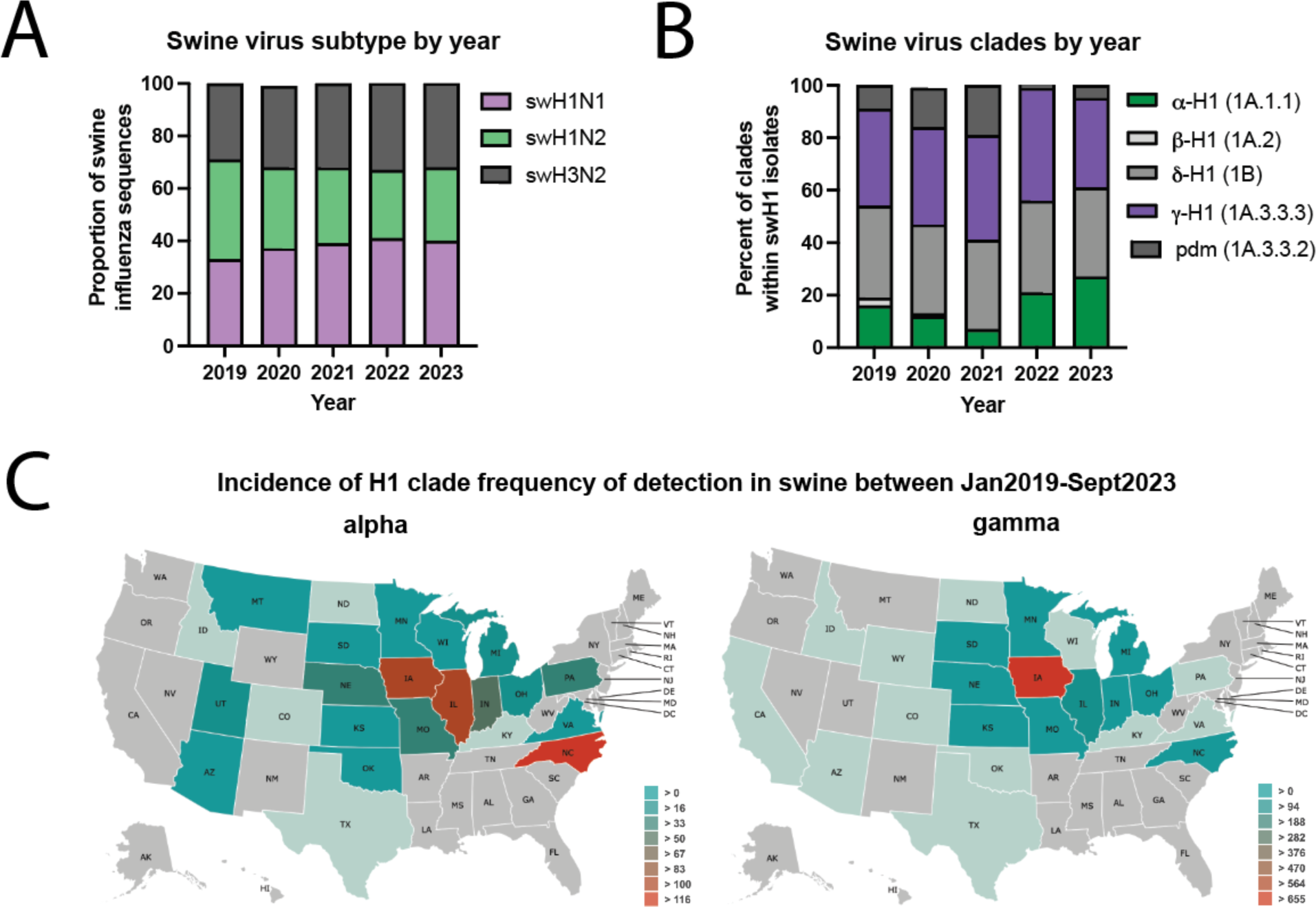
Influenza A virus detected in swine between January 2019 and September 2023 in the USA. **(A)** Influenza A virus subtype detection proportions. **(B)** H1 influenza A virus hemagglutinin clade detection proportions. pdm; pandemic. Data for A and B obtained from octoFLUshow^4^. **(C)** Detections of α-swH1N2 (alpha) and γ -swH1N1 (gamma) influenza A virus in swine across the United States between 2019 and 2021. Data retrieved from ISU FLUture^65^ on September 30, 2023.

**Extended Data Figure 2.**
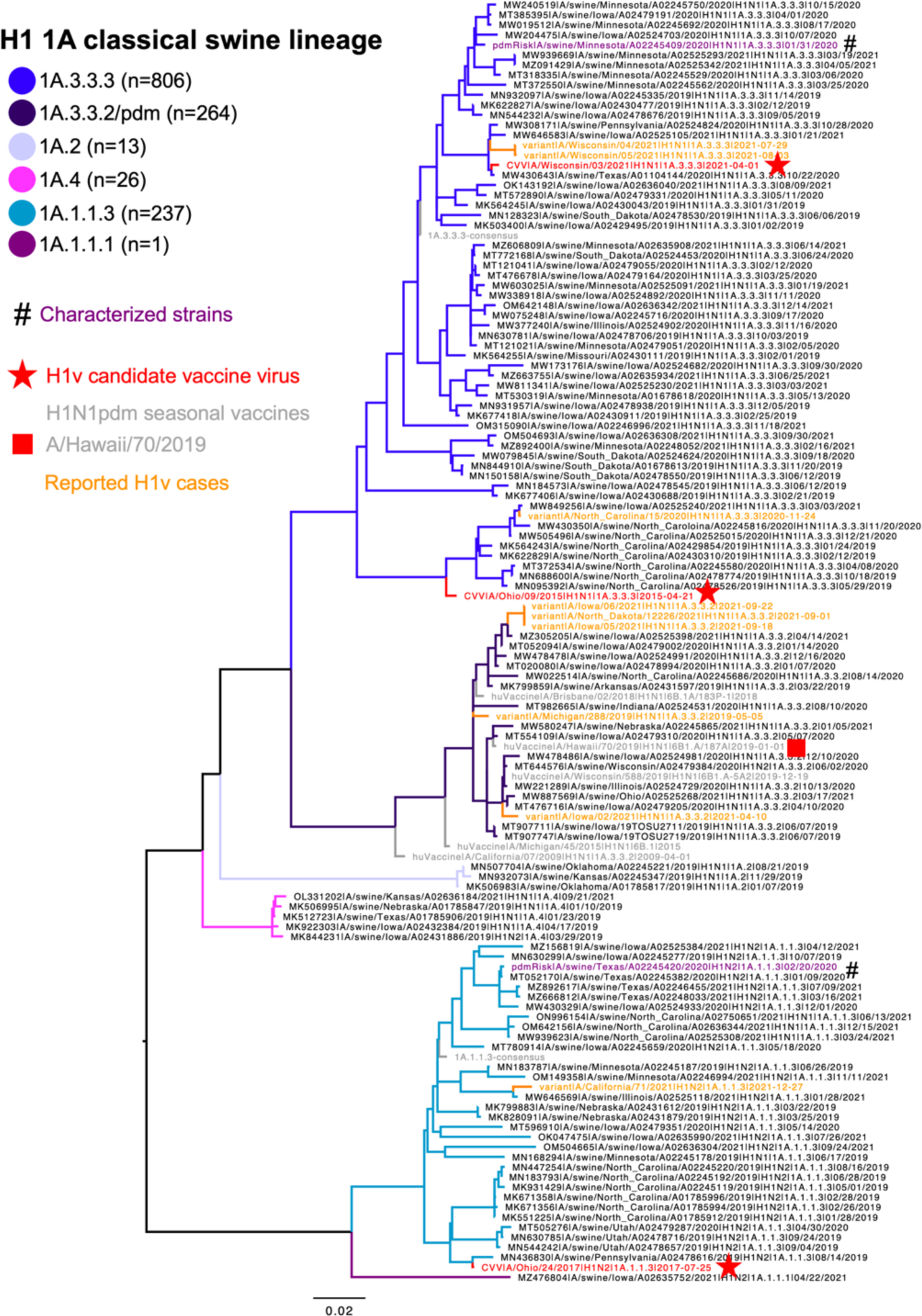
Representative phylogenetic relationships of North American swine H1 1A classical swine lineage influenza A viruses from 2019 to 2021. Each genetic clade was proportionately down sampled using smot^64^ and branches were colored. Swine influenza A virus strains characterized are marked by hash signs (#) and colored purple with the genetic clade consensus colored gray. The numbers in parentheses in the color key indicate number of each genetic clade detected between 2019 and 2021. Human seasonal H1 vaccine strains were colored gray; candidate vaccine viruses were colored red; and reported H1 variant cases detected between 2019 and 2021 were colored orange. The tree was midpoint rooted; all branch lengths are drawn to scale, and the scale bar indicates the number of nucleotide substitutions per site. The complete H1 phylogeny and input data are presented at https://github.com/flu-crew/datasets.

**Extended Data Figure 3.**
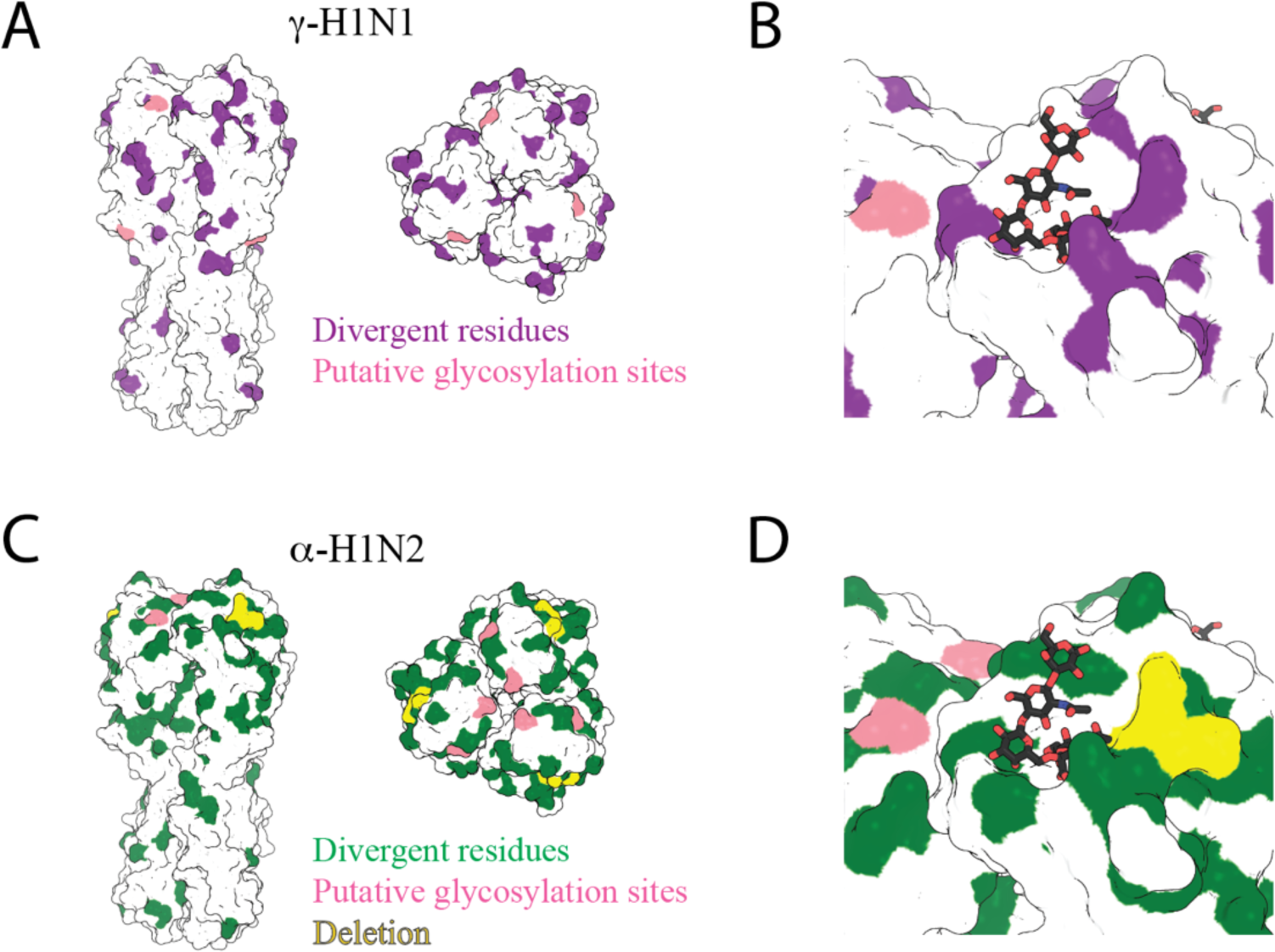
HA structure comparison between H1N1pdm09 and swine viruses. **(A)** Top and side views of a surface representation of HA colored to compare the amino acid sequence of H1N1pdm09 HA and γ-swH1N1. Differences in amino acid sequences are represented in purple and differential putative glycosylation sites are colored pink. **(B)** γ-swH1N1 receptor binding site including α2-6-linked SA (sticks). **(C)** Top and side views of a surface representation of HA comparing the amino acid sequence of H1N1pdm09 HA and α-swH1N2. Differences in amino acid sequences are represented in green and differential putative glycosylation sites are colored pink. The two amino acid residue deletion at residue 133 and 133a (H3N2 numbering) in α-swH1N2 HA are highlighted in yellow. **(D)** α-swH1N2 receptor binding site including α2-6-linked SA (sticks). Images created in PyMOL and is based on PDB 3UBE.

**Extended Data Figure 4.**
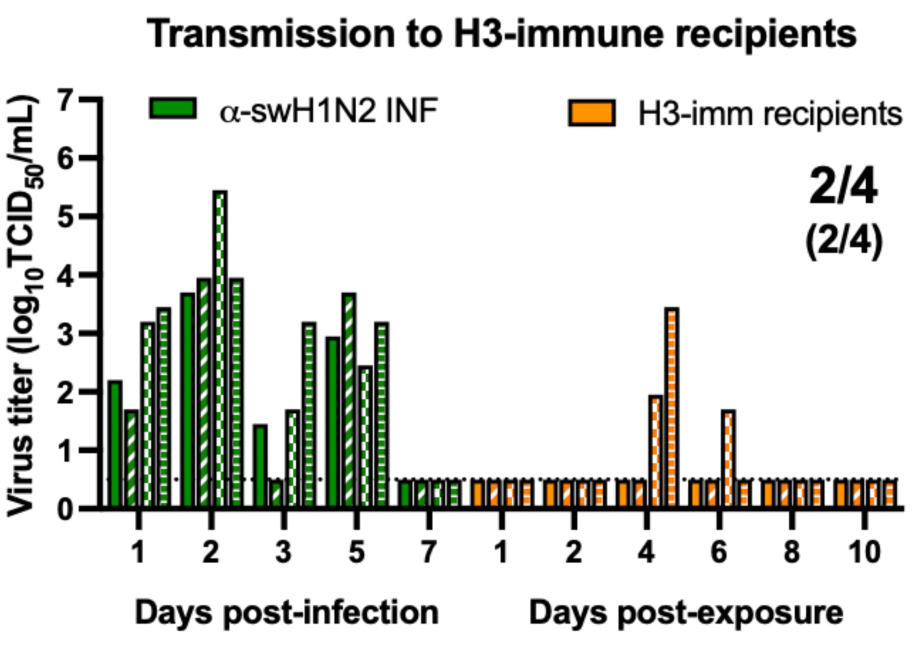
α-swH1N2 transmission to H3N2-imm recipients. Four ferrets were infected with H3N2 A/Perth/16/2009 strain (H3N2-imm) 137 days prior to acting as recipients to α-swH1N2 infected donors. Four donor ferrets were infected with α-swH1N2 and H3N2-imm recipients were placed in the adjacent cage 24 hours later. Nasal washes were collected from all ferrets on the indicated days and titered for virus by TCID_50_. Each bar indicates an individual ferret. For all graphs, the number of recipient ferrets with detectable virus in nasal secretions out of four total is shown; the number of recipient animals that seroconverted at 14- or 21-days post α-swH1N2 exposure out of four total is shown in parentheses. Gray shaded box indicates shedding of the donor during the exposure period. The limit of detection is indicated by the dashed line.

**Extended Data Figure 5.**
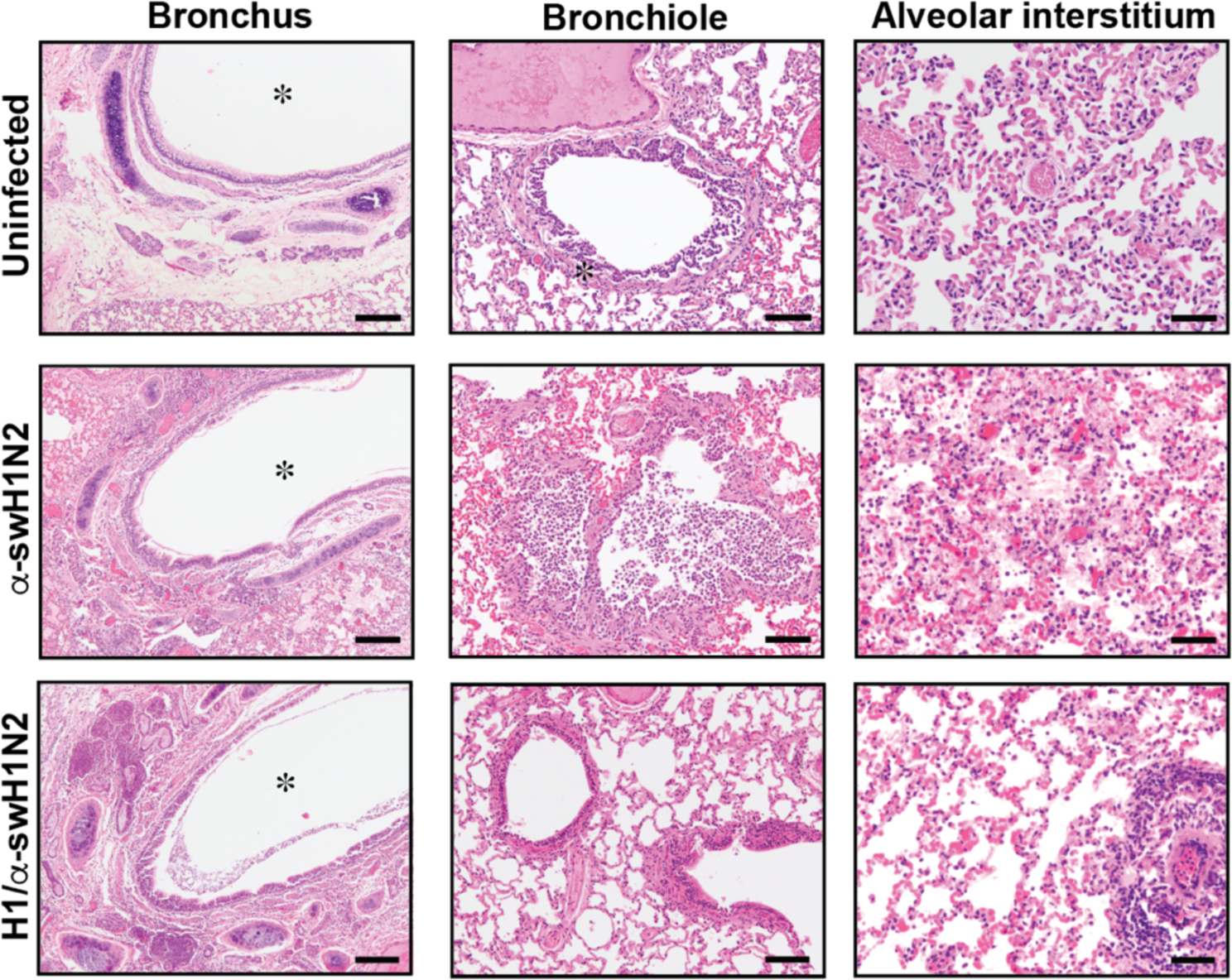
Lung pathology caused by swine α-H1N2 influenza virus is more severe in ferrets with no prior immunity. Lungs from uninfected or α-swH1N2 infected ferrets without or with H1N1pdm09 pre-existing immunity (from Figure 5D) were harvested at 3 dpi. Histopathology was examined by H&E staining. N=2 for each group. Bronchus and bronchiole are at 20x magnification and alveolar interstitium is at 10x. Scale bar is 100 μm for bronchus and bronchiole and 200 μm for alveolar interstitium.

**Extended Data Figure 6.**
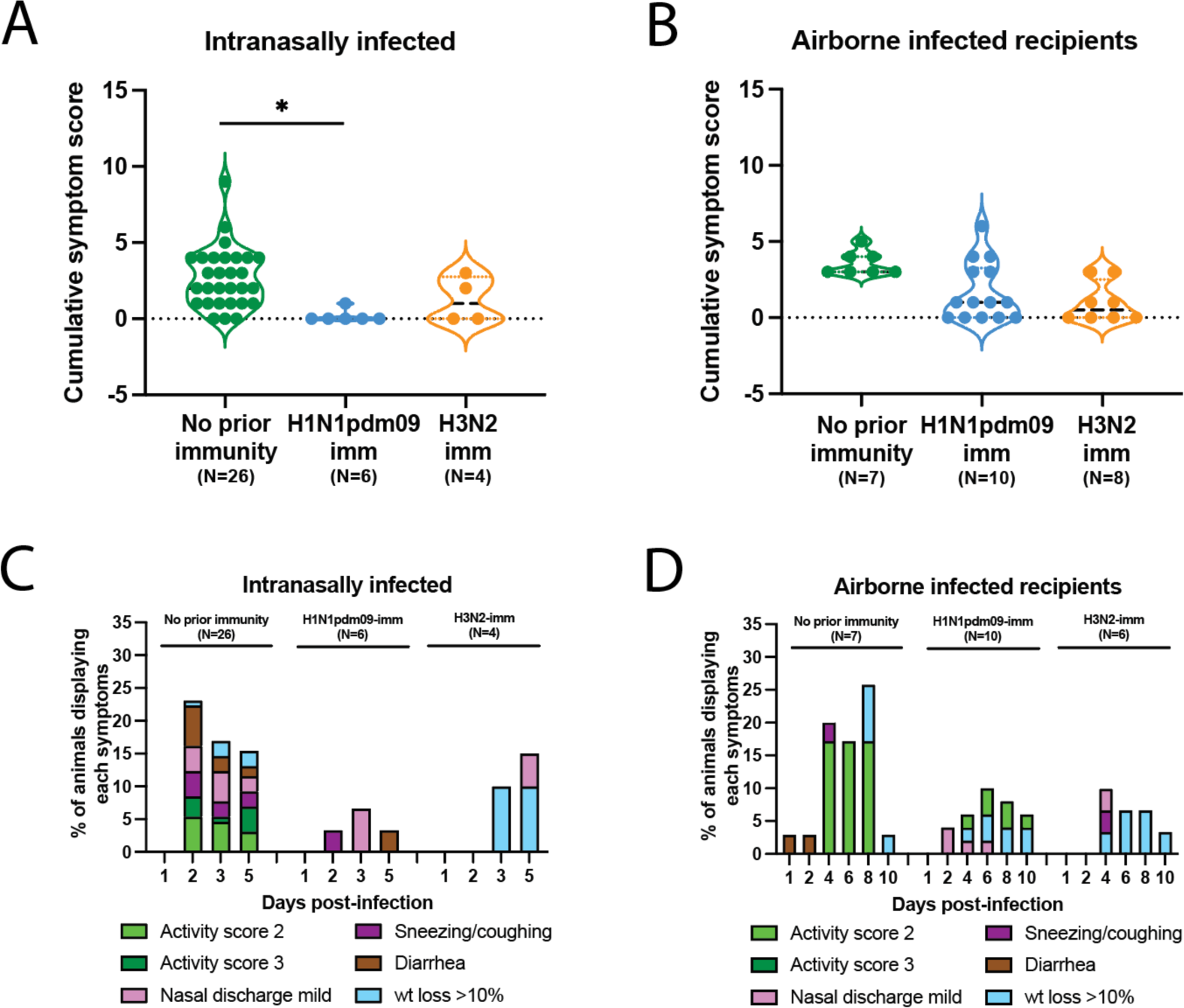
Pre-existing H1N1pdm09 immunity reduces swine α-H1N2 influenza virus clinical signs. **(A)** The symptoms for each intranasally α-swH1N2-infected ferret from Figures 3, 4 and 5 having either no prior immunity (N=22), H1N1pdm09-imm (N=6) or H3N2-imm (N=4) were added together to assign each animal a cumulative score. Each dot represents the cumulative symptoms score for a single ferret. Two-way ANOVA analysis was used to determine statistically significant differences (* p<0.05). **(B)** The symptoms for each airborne α-swH1N2-infected ferret from Figures 3 and 4 having either no prior immunity (N=7), H1N1pdm09-imm (N=10) or H3N2-imm (N=4) were added together to assign each animal a cumulative score. Each dot represents the cumulative symptoms score for a single ferret. **(C)** Percent number of intranasally infected ferrets from panel A displaying each symptom on the indicated days post-infection. **(D)** Percent number of recipient ferrets from panel B displaying each symptom on the indicated days post-exposure.

**Extended Data Figure 7.**
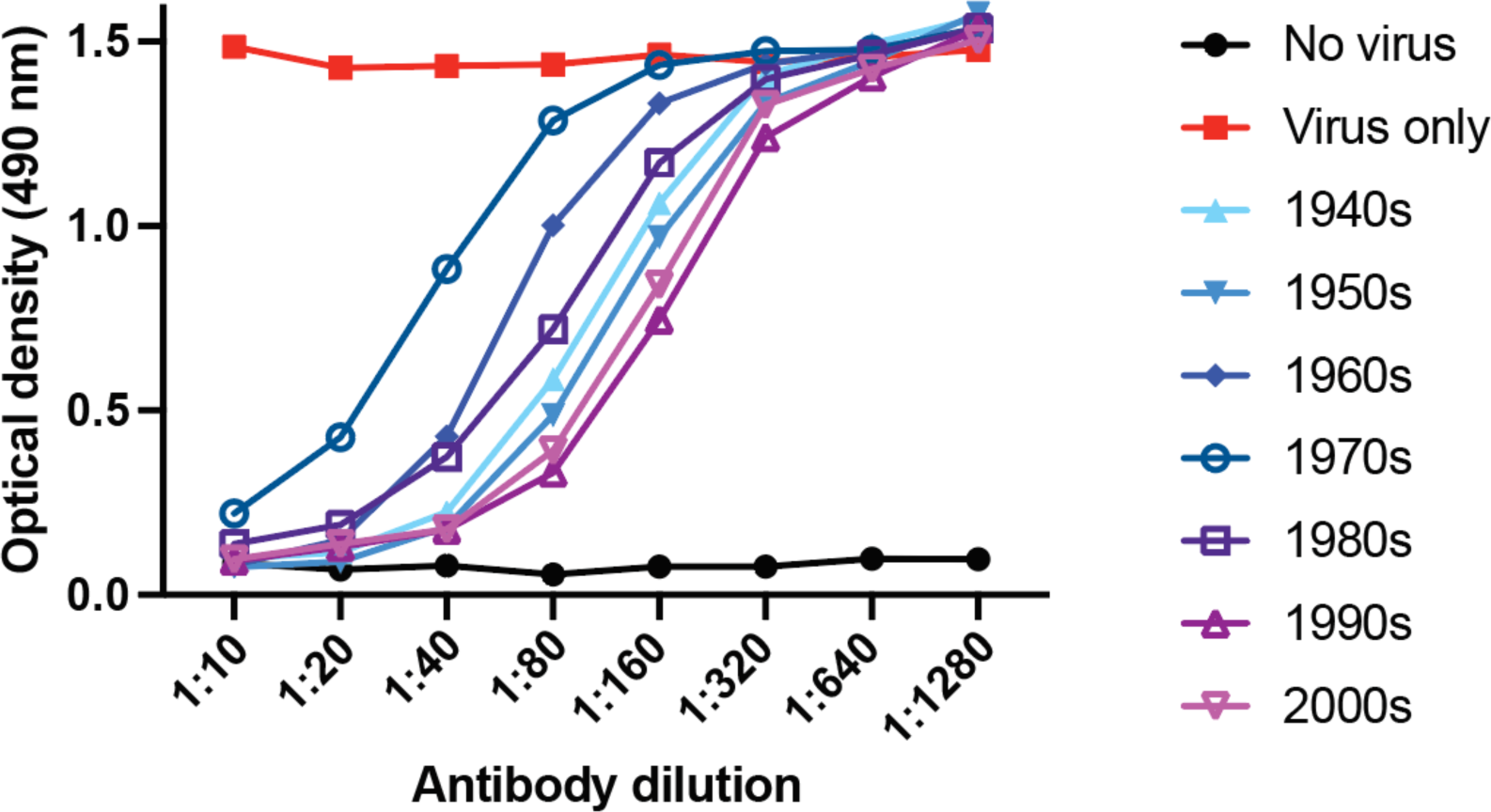
Detection of human anti-NA antibodies that block NA activity. Serially diluted human sera, pooled by birth year, was incubated with an H9 reassortant bearing a NA antigens from A/swine/NY/A01104005/2011. Inhibition of NA activity was tested using fetuin substrate coated plates. Negative control with PBS only (no virus) and positive control with no serum were added as comparators. The data are representative of three experiments run in duplicate.

