## Extended Data Table 1 for "Potential pandemic risk of circulating swine H1N2 influenza viruses"

### Extended Data Table 1: Glycan Structures on Microarray

| | Catalogue | Structure | Symbol |
|---|---|---|---|
| 1 | **M040** | Galβ(1-4)-GlcNAcβ-ethyl-NH2 | 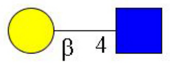 |
| 2 | M221 | | 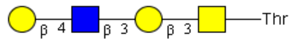 |
| 3 | M222 | | 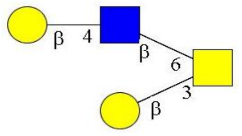 |
| 4 | **M009** | Galβ(1-4)-GlcNAcβ(1-2)-Manα(1-3)-[Galβ(1-4)-GlcNAcβ(1-2)-Manα(1-6)]-Manβ(1-4)-GlcNAcβ(1-4)-GlcNAcβ-Asn-$NH_2$ | 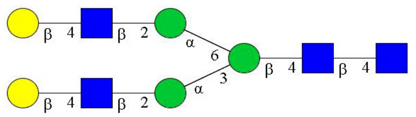 |
| 5 | M226 | | 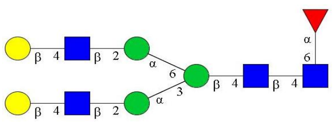 |
| 6 | M227 | | 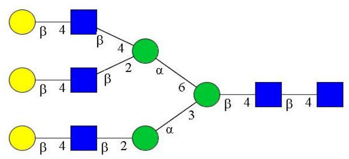 |
| 7 | **SW29** | NeuAca(2-3)-Galβ(1-4)-GlcNAcβ-ethyl-$NH_2$ | 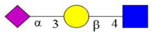 |

| | | | |
|---|---|---|---|
| 8 | **SW30** | NeuAca(2-3)-Galβ(1-4)-GlcNAcβ(1-3)-Galβ(1-4)-GlcNAcβ-ethyl-$NH_2$ | 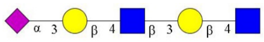 |
| 9 | **SW31** | NeuAcα(2-3)-Galβ(1-4)-GlcNAcβ(1-3)-Galβ(1-4)-GlcNAcβ(1-3)-Galβ(1-4)-GlcNAcβ-ethyl-NH2 | 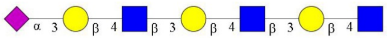 |
| 10 | **M045** | NeuAcα(2-3)-Galβ(1-3)-GalNAcα-Thr-$NH_2$ | 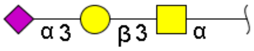 |
| 11 | M120 | 3’ NeuAc LN Core 1 (1163) | 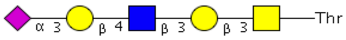 |
| 12 | M128 | 3’ NeuAc DiLN Core 1 (1528) | 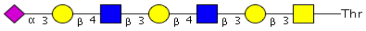 |
| 13 | M142 | 3’ NeuAc TetraLN Core 1 (2259) | 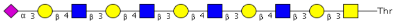 |
| 14 | M143 | 3’ NeuAc PentaLN Core 1 (2624) | 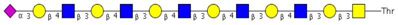 |
| 15 | **M050** | NeuAcα(2-3)-Galβ(1-4)-GlcNAcβ(1-6)-[Galβ(1-3)]-GalNAcα-Thr-$NH_2$ | 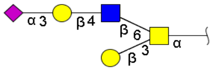 |

| | | | |
|---|---|---|---|
| **16** | **M053** | NeuAcα(2-3)-Galβ(1-4)-GlcNAcβ(1-3)-Galβ(1-4)-GlcNAcβ(1-6)-[Galβ(1-3)]-GalNAcα-Thr-$NH_2$ | 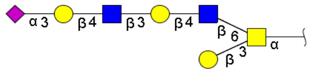 |
| **17** | **M202** | 3’ NeuAc TriLN Core 2 (1894) | 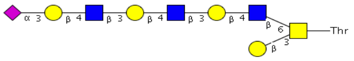 |
| **18** | M152 | 3’ NeuAc TetraLN Core 2 (2259) | 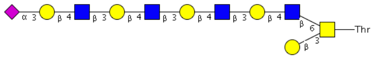 |
| **19** | M149 | 3’ NeuAc PentaLN Core 2 (2624) | 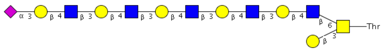 |
| **20** | SW07 | NeuAca2-3Galb1-4GlcNAcb1-2Mana1-3(NeuAca2-3Galb1-4GlcNAcb1-2Mana1-6)Manb1-4GlcNAcb1-4GlcNAc-AsnGly | 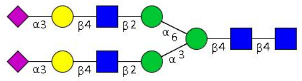 |
| **21** | SW08 | NeuAca2-3Galb1-4GlcNAcb1-3Galb1-4GlcNAcb1-2Mana1-3(NeuAca2-3Galb1-4GlcNAcb1-3Galb1-4GlcNAcb1-2Mana1-6)Manb1-4GlcNAcb1-4GlcNAc-AsnGly | 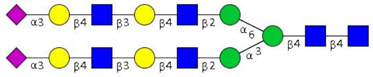 |
| **22** | SW09 | (NeuAca2-3Galb1-4GlcNAcb1-3Galb1-4GlcNAcb1-3Galb1-4GlcNAcb1-2Mana1-6)NeuAca2-3Galb1-4GlcNAcb1-3Galb1-4GlcNAcb1-3Galb1-4GlcNAcb1-2Mana1-3Manb1-4GlcNAcb1-4GlcNAc-AsnGly | 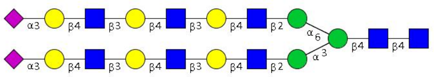 |
| **23** | SW10 | NeuAca2-3Galb1-4GlcNAcb1-2Mana1-3(NeuAca2-3Galb1-4GlcNAcb1-2Mana1-6)Manb1-4GlcNAcb1-4(Fuca1-6)GlcNAc-AsnGly | 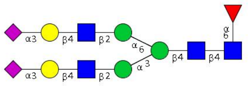 |
| **24** | SW11 | NeuAca2-3Galb1-4GlcNAcb1-3Galb1-4GlcNAcb1-2Mana1-3(NeuAca2-3Galb1-4GlcNAcb1-3Galb1-4GlcNAcb1-2Mana1-6)Manb1-4GlcNAcb1-4(Fuca1-6)GlcNAc-AsnGly | 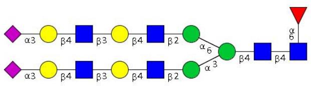 |
| **25** | SW12 | NeuAca2-3Galb1-4GlcNAcb1-3Galb1-4GlcNAcb1-3Galb1-4GlcNAcb1-2Mana1-3(NeuAca2-3Galb1-4GlcNAcb1-3Galb1-4GlcNAcb1-3Galb1-4GlcNAcb1-2Mana1-6)Manb1-4GlcNAcb1-4GlcNAc-AsnGly | 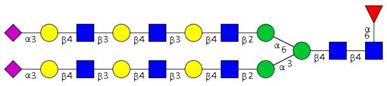 |

| | | | |
|---|---|---|---|
| 26 | **M002** | NeuAca(2-3)-Galβ(1-4)-[Fuca(1-3)]-GlcNAcβ-propyl-$NH_2$ | 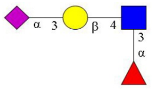 |
| 27 | **M029** | NeuAca(2-3)-Galβ(1-3)-[Fuca(1-4)]-GlcNAcβ(1-3)-Galβ(1-4)-[Fuca(1-3)]-GlcNAcβ-ethyl-$NH_2$ | 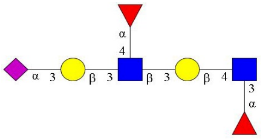 |
| 28 | **M022** | NeuAca(2-3)-Galβ(1-4)-[Fuca(1-3)]-GlcNAcβ(1-3)-Galβ(1-4)-[Fuca(1-3)]-GlcNAcβ-ethyl-$NH_2$ | 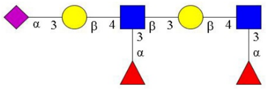 |
| 29 | **M015** | NeuAca(2-3)-Galβ(1-4)-[Fuca(1-3)]-GlcNAcβ(1-3)-Galβ(1-4)-[Fuca(1-3)]-GlcNAcβ(1-3)-Galβ(1-4)-[Fuca(1-3)]-GlcNAcβ-ethyl-$NH_2$ | 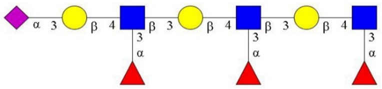 |
| 30 | **M206** | 3' SLeX TriLN Core 1(2332) | 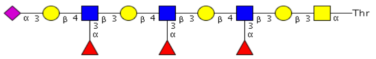  |
| 31 | M147 | 3' SLeX TriLN Core 3(2170) | 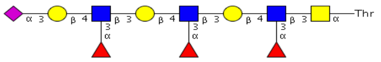  |
| 32 | M215 | NeuAc(2-6)-Galb(1-4)-(6S)GlcNacb-ethyl-NH2 | 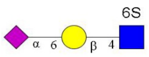  |
| 33 | **M003** | NeuAca(2-6)-Galβ(1-4)-6-O-sulfo-GlcNAcβ-propyl-$NH_2$ | 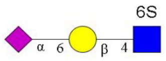  |

| | | | |
|---|---|---|---|
| **34** | **SW32** | NeuAca(2-6)-Galβ(1-4)-GlcNAcβ-ethyl-$NH_2$ | 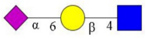 |
| **35** | **SW33** | NeuAca(2-6)-Galβ(1-4)-GlcNAcβ(1-3)-Galβ(1-4)-GlcNAcβ-ethyl-$NH_2$ | 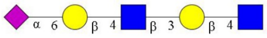 |
| **36** | **SW34** | NeuAca(2-6)-Galβ(1-4)-GlcNAcβ(1-3)-Galβ(1-4)-GlcNAcβ(1-3)-Galβ(1-4)-GlcNAcβ-ethyl-$NH_2$ | 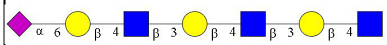 |
| **37** | M121 | 6' NeuAc LN Core 1 (1163) | 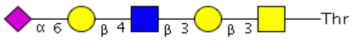 |
| **38** | M129 | 6' NeuAc DiLN Core 1 (1528) | 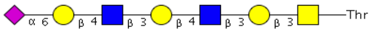 |
| **39** | M154 | 6' NeuAc TriLN Core 1 (1894) | 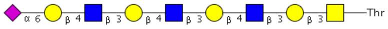 |
| **40** | **M051** | NeuAcα(2-6)-Galβ(1-4)-GlcNAcβ(1-6)-[Galβ(1-3)]-GalNAcα-Thr-$NH_2$ | 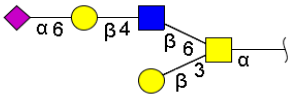 |
| **41** | **M054** | NeuAcα(2-6)-Galβ(1-4)-GlcNAcβ(1-3)-Galβ(1-4)-GlcNAcβ(1-6)-[Galβ(1-3)]-GalNAcα-Thr-$NH_2$ | 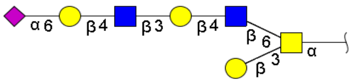 |

| | | | |
|---|---|---|---|
| **42** | **M201** | 6’ NeuAc TriLN Core 2 (1894) | 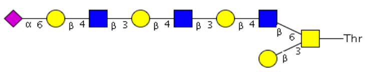 |
| **43** | M159 | 6’ NeuAc TetraLN Core 2 (2259) | 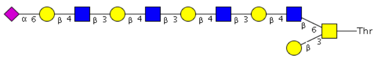 |
| **44** | M157 | 6’ NeuAc PentaLN Core 2 (2624) | 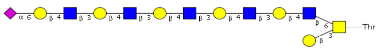 |
| **45** | SW01 | NeuAca2-6Galb1-4GlcNAcb1-2Mana1-3(NeuAca2-6Galb1-4GlcNAcb1-2Mana1-6)Manb1-4GlcNAcb1-4GlcNAc-AsnGly | 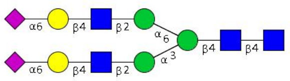 |
| **46** | SW02 | NeuAca2-6Galb1-4GlcNAcb1-3Galb1-4GlcNAcb1-2Mana1-3(NeuAca2-6Galb1-4GlcNAcb1-3Galb1-4GlcNAcb1-2Mana1-6)Manb1-4GlcNAcb1-4GlcNAc-AsnGly | 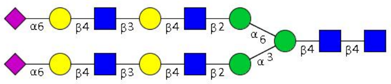 |
| **47** | SW03 | NeuAca2-6Galb1-4GlcNAcb1-3Galb1-4GlcNAcb1-3Galb1-4GlcNAcb1-2Mana1-3(NeuAca2-6Galb1-4GlcNAcb1-3Galb1-4GlcNAcb1-3Galb1-4GlcNAcb1-2Mana1-6)Manb1-4GlcNAcb1-4GlcNAc-AsnGly | 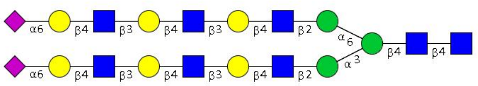 |
| **48** | SW04 | NeuAca2-6Galb1-4GlcNAcb1-2Mana1-3(NeuAca2-6Galb1-4GlcNAcb1-2Mana1-6)Manb1-4GlcNAcb1-4(Fuca1-6)GlcNAc-AsnGly | 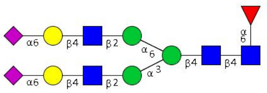 |
| **49** | SW05 | NeuAca2-6Galb1-4GlcNAcb1-3Galb1-4GlcNAcb1-2Mana1-3(NeuAca2-6Galb1-4GlcNAcb1-3Galb1-4GlcNAcb1-2Mana1-6)Manb1-4GlcNAcb1-4(Fuca1-6)GlcNAc-AsnGly | 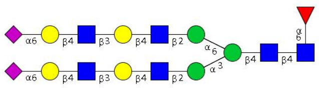 |
| **50** | SW06 | NeuAca2-6Galb1-4GlcNAcb1-3Galb1-4GlcNAcb1-3Galb1-4GlcNAcb1-2Mana1-3(NeuAca2-6Galb1-4GlcNAcb1-3Galb1-4GlcNAcb1-3Galb1-4GlcNAcb1-2Mana1-6)Manb1-4GlcNAcb1-4GlcNAc-AsnGly | 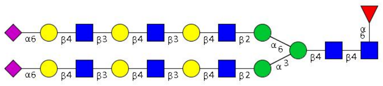 |
